# Dissociable Cellular and Genetic Mechanisms of Cortical Thinning at Different Life Stages

**DOI:** 10.1101/2022.03.21.484899

**Authors:** Amirhossein Modabbernia, Didac Vidal-Pineiro, Ingrid Agartz, Ole A Andreassen, Rosa Ayesa-Arriola, Alessandro Bertolino, Dorret I Boomsma, Josiane Bourque, Alan Breier, Henry Brodaty, Rachel M Brouwer, Jan K Buitelaar, Erick J Canales-Rodríguez, Xavier Caseras, Patricia J Conrod, Benedicto Crespo-Facorro, Fabrice Crivello, Eveline A Crone, Greig I de Zubicaray, Erin W Dickie, Danai Dima, Stefan Frenzel, Simon E Fisher, Barbara Franke, David C Glahn, Hans-Jörgen Grabe, Dominik Grotegerd, Oliver Gruber, Amalia Guerrero-Pedraza, Raquel E Gur, Ruben C Gur, Catharina A Hartman, Pieter J Hoekstra, Hilleke E Hulshoff Pol, Neda Jahanshad, Terry L Jernigan, Jiyang Jiang, Andrew J Kalnin, Nicole A Kochan, Bernard Mazoyer, Brenna C McDonald, Katie L McMahon, Lars Nyberg, Jaap Oosterlaan, Edith Pomarol-Clotet, Joaquim Radua, Perminder S Sachdev, Theodore D Satterthwaite, Raymond Salvador, Salvador Sarro, Andrew J Saykin, Gunter Schumann, Jordan W Smoller, Iris E Sommer, Thomas Espeseth, Sophia I Thomopoulos, Julian N Trollor, Dennis van ‘t Ent, Aristotle Voineskos, Yang Wang, Bernd Weber, Lars T Westlye, Heather C Whalley, Steven CR Williams, Katharina Wittfeld, Margaret J Wright, Paul M Thompson, Thomas Paus, Sophia Frangou

## Abstract

Mechanisms underpinning age-related variations in cortical thickness in the human brain remain poorly understood. We investigated whether inter-regional age-related variations in cortical thinning (in a multicohort neuroimaging dataset from the ENIGMA Lifespan Working Group totalling 14,248 individuals, aged 4-89 years) depended on cell-specific marker gene expression levels. We found differences amidst early-life (<20 years), mid-life (20-60 years), and late-life (>60 years) in the patterns of association between inter-regional profiles of cortical thickness and expression profiles of marker genes for CA1 and S1 pyramidal cells, astrocytes, and microglia. Gene ontology and enrichment analyses indicated that each of the three life-stages was associated with different biological processes and cellular components: synaptic modeling in early life, neurotransmission in mid-life, and neurodegeneration in late-life. These findings provide mechanistic insights into age-related cortical thinning during typical development and aging.

## Introduction

The human cerebral cortex undergoes substantial thinning, beginning in late childhood and extending throughout the lifespan^1,2^. These macro-scale changes are underpinned by cellular and molecular processes that vary across life-stages. Cortical thinning during development is likely attributable to adaptive synaptic and dendrite elimination ^3–5^ combined with increased intracortical myelin ^6,7^. In late adulthood it has been attributed to deterioration in vasculature, shrinkage of large neurons ^8^, loss of myelinated axonal fibers ^9^, and regressive synaptic density reduction ^10^. Elucidating the cellular and genetic underpinning of the typical trajectory of cortical thinning provides the foundation for understanding how deviations may occur in development and ageing.

To address this challenge, our group developed a virtual-histology approach that is based on the spatial correlation – across the human cerebral cortex – of gene-expression and magnetic resonance imaging (MRI)-derived data ^11^. In our prior studies using this approach, we found that higher expression of marker genes of CA1 pyramidal cells, astrocytes, and microglia was associated with less cortical thinning in childhood and adolescence ^12,13^, and with accelerated cortical thinning in late-life^13^. These findings suggest that biological mechanisms supported by the same cellular populations may differ according to life-stage. Higher expression of marker genes for CA1 pyramidal cell may contribute to neurite sprouting during development ^14^ and shrinkage during ageing ^15^, while higher expression of marker genes for glial cells may support the consolidation of brain organization in early life ^16^, but facilitate a heightened proinflammatory profile in the ageing brain ^17^.

In the present study, we extended these investigations in two distinct directions. First, we used one of the largest available samples of neuroimaging data from healthy participants (N=14,248) covering the human lifespan from the Enhancing Neuroimaging Genetics through Meta-analysis (ENIGMA) consortium ^18^. Leveraging these data, our first aim was to test the reproducibility of the prior findings regarding the association of cell-specific gene expression and cortical thinning, which have relied on smaller samples (<4,000). Our second aim was to use gene co-expression and over-representation analyses to test whether the cell-specific gene sets associated with cortical thinning implicate differing biological processes and cellular components that support either developmental or regressive events at different life-stages.

## Materials and Methods

### Neuroimaging Dataset

De-identified cross-sectional demographic data and MRI-derived estimates of cortical thickness from 35 cohorts participating in the ENIGMA Lifespan Working Group were pooled to create the neuroimaging dataset. The pooled sample comprised 14,248 individuals (52.7% female) aged 4-89 years. Ethical approval and informed consent were overseen by the investigators of each cohort, and data were shared in accordance with the ENIGMA data-use agreements and local policies. Participants in each cohort had been screened to exclude psychiatric disorders, medical and neurological morbidity, and cognitive impairment. Details of demographic characteristics and screening procedures are presented in Supplementary Tables 1 and 2 and Supplementary Figure 1.

### Estimation of age-related cortical thinning

Standardized ENIGMA analysis and quality assurance protocols were applied to T_1_-weighted whole-brain MRI scans to extract estimates of thickness from cortical regions defined by the Desikan-Killiany Atlas ^19^ based on the FreeSurfer pipelines (http://surfer.nmr.mgh.harvard.edu). Details of the neuroimaging analyses and quality assurance protocols are presented in supplemental data and Supplemental Table 2. Only the left-hemisphere measures (N=34) were used in subsequent analyses to align with the availability of the gene-expression data. Separate generalized additive mixed models (GAMM; implemented with the *mgcv* R-package) ^20^ were used to model the association between age and each of the 34 regional measures of cortical thickness, adjusted for sex, as fixed effects, and scanner identifiers as random intercepts. The GAMM does not assume linearity in the relationship between outcome and predictors and uses flexible functions (i.e., splines) to model nonlinear relationships. Splines behave as polynomial functions, each covering a small range of the data, which converge on each other at different points (i.e., knots). For each region, we specified cubic splines as functional approximators and evaluated models with an increasing number of knots beginning at 5 (see also sensitivity analyses in Supplementary Table 3 and Supplementary Figure 3). The model that used one degree of freedom less than what was allowed was considered optimal, allowing us to tailor the model parameters to each region while minimizing overfitting. We then employed the *gratia* R-package ^21^, which uses a finite differences approach, to compute the GAMM derivatives as an estimate of cortical thinning of each region for every 1-year shift in chronological age. As the age range of the ENIGMA Lifespan dataset is 4-89 years, these analyses resulted in 86 agespecific inter-regional profiles of cortical thinning across the 34 cortical regions (Supplementary Figure 2). Each age-specific inter-regional profile represents the spatial variation in cortical thinning across the 34 cortical regions, at that specific age.

### Cell-specific gene expression profiling

Cell-specific gene expression profiling followed our previously developed methods ^11–13,22^ using the Allen Human Brain Atlas (AHBA) ^23^ as the source of gene expression data (details in the Supplement section A4). Subsequent analyses were restricted to a panel of 2,511 genes, henceforth referred to as the *consistent genes panel*, identified by the consistency of their interregional expression in the 34 left-hemisphere cortical regions in the AHBA (across donors) and between AHBA and the BrainSpan dataset. ^12^. The interregional profile of gene expression represents the spatial variation in the expression of a given gene across these cortical regions. Cell-specific gene labels provided by Zeisel and colleagues (2015) were updated as per Mancarci and French ^24^ and clustered into 9 cell-specific gene panels: S1 pyramidal neurons (n□=□73 human gene symbols), CA1 pyramidal neurons (n□=□103), interneurons (n□=□100), astrocytes (n□=□54), microglia (n□=□48), oligodendrocytes (n□=□60), ependymal (n□=□84), endothelial (n□=□57), and mural (pericytes and vascular smooth muscle cells; n□=□25).

### Study-Specific Gene Expression Database and Co-Expression Networks

We created a study-specific gene-expression database based on information from post-mortem cortical tissue from 572 unique donors, aged 0 to 102 years at the time of death, by harmonizing data from five databases: the AHBA, the BrainCloud ^25,26^, the Brain eQTL Almanac project ^27^, the BrainSpan (http://brainspan.org), and the Genotype-Tissue Expression Project ^28^. The study-specific database included expression levels for 15,380 genes that were available in all five databases, 2,321 of which were also part of the consistent genes panel. The co-expression matrix of each of these 2,321 genes was determined using linear mixed models and ranked according to effect size (details in Supplement Section 5A).

### Statistical Analyses

Statistical procedures throughout the study were implemented in R (www.r-project.org, v. 3.6.1) (Team RC, 2013). The threshold for statistical significance was adjusted for multiple testing using false discovery rate (FDR) correction.

#### Correlation between lifespan cortical thinning and cell-specific gene expression profiles

We quantified associations between each inter-regional profile of cortical thinning (n=86; obtained from the GAMM derivative sampled across the entire age-range of the sample) and interregional expression of each cell-specific gene (N=604) using the Pearson’s correlation coefficient as per our previous work ^12,13^. We computed the Pearson’s correlation between the thinning and the gene expression profiles, for each cortical profile (n = 86, one profile for each age from 4 to 89 years) and consistent cell-specific gene panels (n =9). The statistical significance of these associations was established through resampling and permutation testing against a null distribution generated for each actual cell-specific gene panel, by assessing the correlation between cortical-thinning profiles and pseudo-panels of randomly selected cell-specific genes of equal number to each of the actual panels. Statistical significance was established by combining Bonferroni-like correction for the number of inter-regional cortical thinning profiles (n=86) and FDR correction for the number of cell-specific panels (details in supplementary data).

#### Gene Co-Expression and Enrichment

Gene co-expression networks were generated by considering all the consistent genes for which information was available in the study-specific database (N=2,311), irrespective of their cell-specific assignment, as several of these genes are likely to influence biological pathways across multiple cell types. The co-expression matrix of these 2,321 genes was determined using linear mixed models and ranked according to effect size (details in supplemental data). Guided by the preceding analyses, we selected those interregional cortical thinning profiles that had significant correlations with inter-regional cell-specific gene expression profiles; these inter-regional cortical-thinning profiles were then averaged to generate a single life-stage inter-regional cortical-thinning profile for three paradigmatic age-groups consistent with early life (below 20 years of age), mid-life (20-60 years of age) and late-life (over 60 years of age). Quantification of associations between each life-stage cortical thinning profile and gene co-expression networks proceeded as per Sliz and colleagues ^29^. Accordingly, we indexed those cell-specific genes showing high fidelity, i.e., genes that were in the top 5% among the genes consistently associated with the cortical-thinning profile. Based on our previous work^22^, we selected the top 0.1% of the genes in the study-specific co-expression database that were co-expressed with the high-fidelity subset of the consistent genes panel. We then conducted gene ontology (GO) enrichment analysis using the *clusterProfiler* package ^30^. GO provides a unified vocabulary to describe gene functions and their inter-relations in terms corresponding to biological processes (BP) and cellular components (CC). BP and CC gene sets between 10 and 500 genes were considered. The threshold of statistical significance was set at *P_FDR_*<0.01 to account for the three life-stages. The enrichment analysis was generally invariant to the number of genes most co-expressed with the high-fidelity genes, as well as celltype and age-range-specific analyses (details in Supplement Section 6A).

## Results

### Correlation between lifespan cortical thinning and cell-specific gene expression profiles

The MRI-based lifespan changes in regional cortical thickness displayed the prototypic pattern of decline (Supplementary Figure 2). Several inter-regional cell-specific gene profiles showed significant correlations with the inter-regional cortical thinning profiles (Figure 1 and 2) that were robust to the GAMM parameters for cortical thickness derivative estimates (Supplementary Figure 3). Positive correlation coefficients indicate less cortical thinning with higher expression levels of a cell-specific gene profile and the opposite is the case for negative coefficients. Positive correlations between inter-regional cortical thinning profiles were observed with the inter-regional expression profiles of astrocyte-specific genes both in early-life (below 16 years of age) and in mid-life (between the ages of 45-60 years). Negative correlations between interregional cortical thinning and interregional expression levels of S1-pyramidal-specific genes were found both in early-life (below 23 years of age) and in mid-life (between the ages of 36-64 years). Correlations between inter-regional cortical thinning profiles with inter-regional expression levels of microglia- and CA1 pyramidal-specific genes were positive in early-life (below 18 years of age) but negative in late-life (after the age of 67 years). Inter-regional expression levels of ependymal-specific and oligodendrocyte-specific genes respectively showed positive and negative correlations with inter-regional cortical thinning profiles in mid-life (between 45-60 years of age).

**Figure 1.**
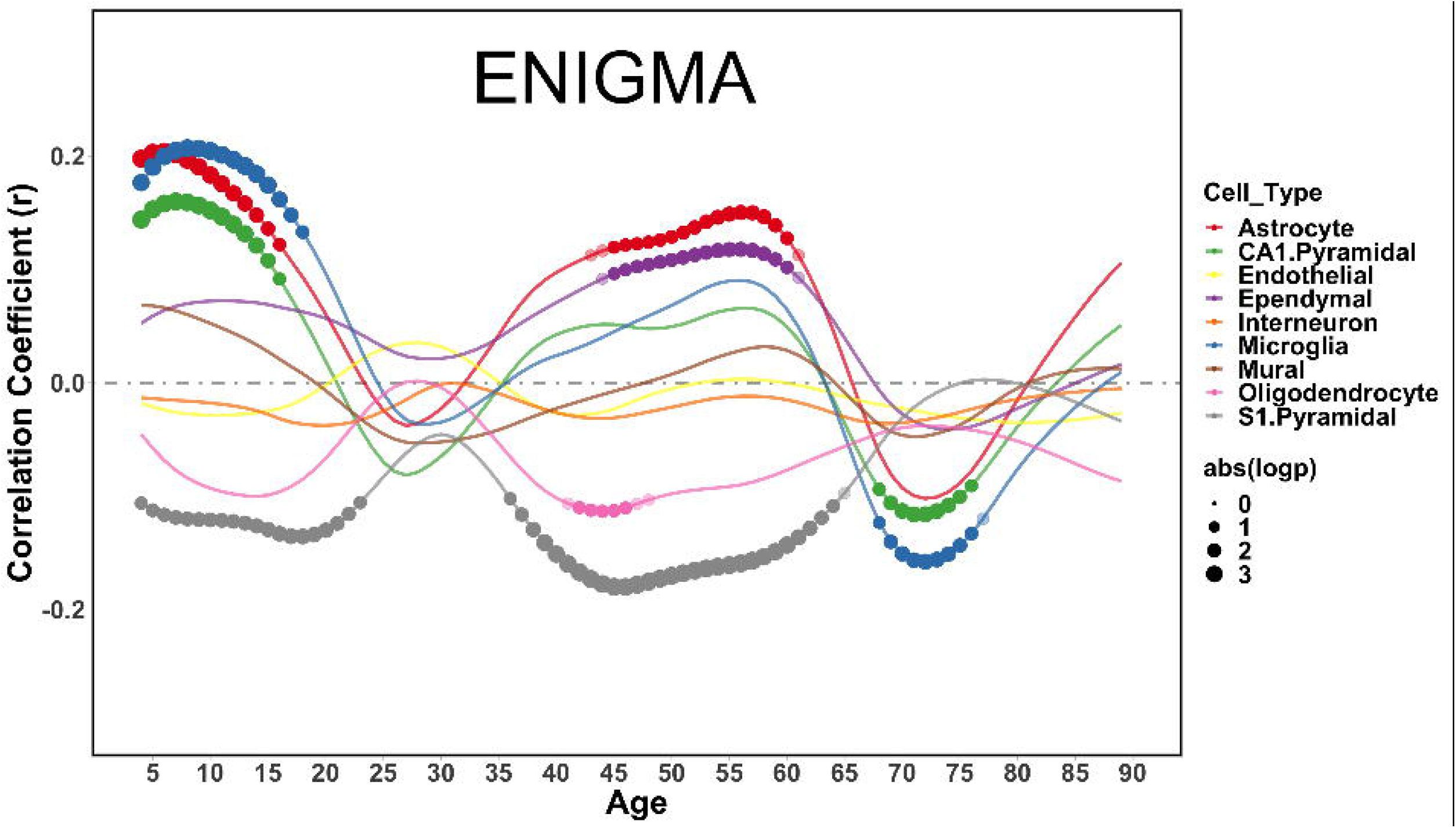
Virtual Histology of Cortical Thinning Profiles Across the Lifespan. Significant correlations between inter-regional cortical thinning and cell-specific gene expression profiles across the lifespan. The horizontal axis represents age in years and the vertical axis the correlation coefficients. Positive correlation coefficients indicate less cortical thinning with higher expression levels of a cell-specific gene profile and the opposite is the case for negative coefficients. Color key: Astrocytes=Red; Mural=Brown; CA1 Pyramidal Cells=Green; Endolthelial=Yellow; Ependymal Cells=Purple; Interneuron=Orange; Microglia=Blue; Oligodendrocytes=Pink; S1 Pyramidal Cells=Gray.

**Figure 2.**
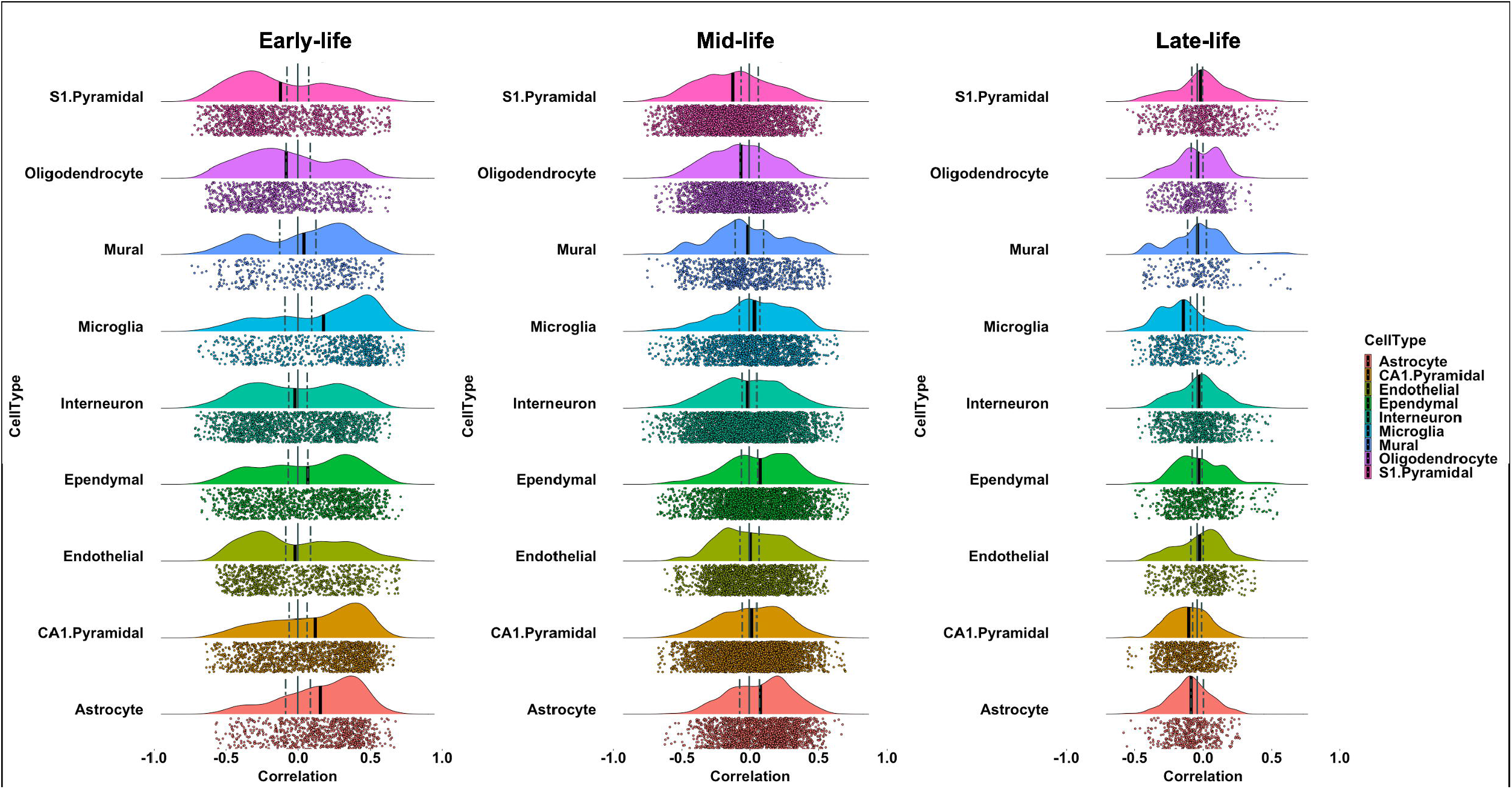
Distribution of the Correlation Coefficients of Between Life-Stage Cortical Thickness Profiles and Inter-Regional Cell-Specific Gene Expression Levels in Early-, Mid- and Late-Life. Each plot presents the distribution of the correlation coefficients of between life-stage cortical thickness profiles and the average inter-regional gene expression levels for each type of cells, in early-life (<20 years of age), mid-life (20-60 years of age), and late-life (>60 years of age). The horizontal axis represents the correlation coefficients between cortical thinning and expression profiles for each set of cell-specific marker genes and the vertical axis indicates the estimated probability density for the correlation coefficients. Each dot represents a correlation value between cortical thinning for a given age (in the corresponding age group) and cortical gene expression pattern related to any one of the cell-type specific genes (for the corresponding celltype). In each age group, the widest age range with FDR-corrected significant results is chosen to generate the figure. The solid black line represents the mean correlation coefficient between the cortical thinning profile and gene-expression for that specific cell-type in that age-group. The dashed lines represent the mean and 95% confidence intervals for the null distribution.

### Association of life-stage inter-cortical thinning profiles with gene co-expression and enrichment

Significant associations of the early-life inter-regional cortical thinning profile with GO:BP terms related to the regulation of trans-synaptic signaling, synaptic pruning, and cognition (Figure 3, Supplementary Figure 4) while associations with GO:CC terms related to synaptic and post-synaptic structures, glutamatergic transmission, and secretory granules membrane, a component of microglial cells (Figure 3, Supplementary Figure 5). Significant associations of the mid-life-stage inter-regional cortical thinning profile with GO: BP terms related to neurotransmitter, amino-acid, and synaptic vesicle transport (Figure 3, Supplementary Figure 6) while associations with GO:CC terms related to synaptic and vesicular membrane components and glutamatergic synapses (Figure 3, Supplementary Figure 7). Significant associations of the late-life-stage inter-cortical thinning profile with GO:BP terms related to the Ras protein signal transduction and acute inflammatory responses (Figure 3, Supplementary Figure 8), while GO:CC terms related to exocytic and synaptic vesicles and components of synapses involved in γ-aminobutyric acid (GABA) and glutamate function (Figure 3, Supplementary Figure 9). These results were robust to selecting genes related to a specific cell-type, selecting the exact agerange where the gene-expression cortical thinning association was significant for a specific celltype, and different thresholding parameters for the most significant genes and co-expression patterns (Online Figures-see Supplement section B4)

**Figure 3.**
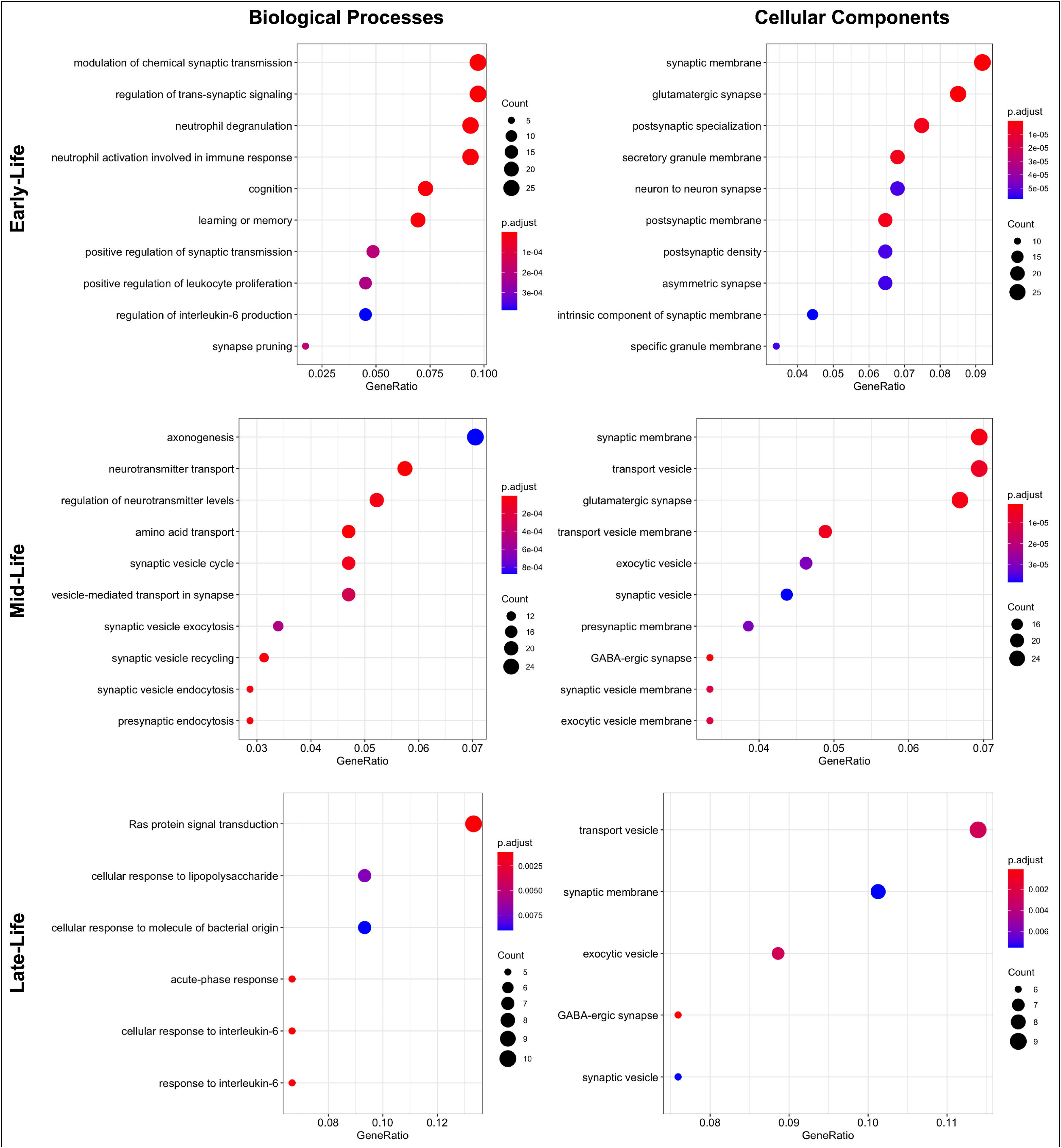
Gene Ontology and Enrichment Analysis for Biological Processes and Cellular Components for Early-, Mid- and Late-Life. Each panel represents the number of genes corresponding to gene ontology terms for biological processes and cellular components that were significantly associated with cortical thinning in early-life (<20 years of age), mid-life (20-60 years of age), and late-life (>60 years of age). The size of each circle represents the gene count and its color the *P*-value of the enrichment analysis results.

## Discussion

Using MRI-based estimates of cortical thickness obtained from a multicohort sample of 14,248 healthy individuals, aged 4-89 years, we identified distinct spatial patterns of cell-specific gene expression, biological processes and cellular components underlying cortical thinning at different life-stages across the human cerebral cortex.

Higher expression levels of marker genes for microglia and CA1 pyramidal cells were associated with less cortical thinning in early life (<20 years) but with accelerated cortical thinning in late life (>60 years). These results largely replicate our previous findings in an adolescent sample ^12^, and in a smaller neuroimaging sample (N=4,004) with lifespan coverage ^13^. The opposing associations between cortical thinning and expression levels of marker genes for CA1 pyramidal cells in early and late-life are also aligned with our previous findings ^12,13^, as well as with histological findings of neurite sprouting during development and shrinkage during ageing ^14,15^. The same pattern was also noted for the expression of marker genes for microglia; this is consistent with their role in adaptive elimination of neurites and synapses during development and in facilitating pro-inflammatory responses in ageing ^16,17^. The current study also confirmed the association between the level of expression of marker genes for S1 pyramidal cells and accelerated cortical thinning in early-life ^12^, and suggests that this association persists in mid-life. Marker genes for the S1 pyramidal cells are known to include those that regulate potassium signaling ^12^, which has been linked to cortical thinning during the consolidation of brain organization in development ^31^ and through apoptosis later in life ^30,32^. We did not detect significant associations between cortical thinning and expression levels of gene markers for oligodendrocytes during early life, which is partly consistent with prior studies ^12,13^. Nonetheless, accelerated cortical thinning has been associated with higher oligodendrocyte-specific genes in male adolescents ^12^, and with higher expression levels of a myelin-specific gene panel in children and adolescents regardless of sex ^33^.

The gene co-expression and enrichment supported and extended the virtual histology results. Across three paradigmatic age-groups, corresponding to early-, mid- and late-life, genes with the most substantial association with the corresponding life-stage cortical thinning profile and their co-expression networks were identified and tested for enrichment. In early life, enrichment identified genes involving GO:BP and GO:CC terms relating to cognition and the formation and maturation of synaptic circuits. In mid-life, enrichment involved GO:BP terms relating to neurotransmitter synthesis, transport and secretion and GO:CC terms, which support these processes across neurotransmitter systems in general, and for glutamatergic and GABAergic synapses in particular. Neurotransmission is the most important process of neuronal communication, and this pattern of enrichment highlights its importance for cortical organization throughout mid-life. In late life, GO:BP and GO:CC terms relating to Ras protein transduction and inflammation were most prominent. The human Ras superfamily of small guanosine triphosphatases (GTPases) has over 150 members implicated in multiple biological pathways ^34^. Some of these RAS protein transduction genes, such as the CDC42SE2, the CDH13, and the NKAP1A have been implicated in the aetiology of neurodegenerative diseases ^35–38^. Enrichment in GO:BP terms such as the cellular response to lipopolysaccharide in late, but not early life, supports the notion of a pro-inflammatory function for microglia in brain ageing ^39^.

An inherent limitation of the virtual histology approach is that the neuroimaging and the cellspecific gene expression data originate with different individuals. We have already demonstrated, however, the value of this approach in yielding mechanistic insights with regards to typical age-related cortical thinning ^12,13,33,40^, and cortical reductions associated with psychiatric disorders ^13,22^. The current study benefits from a large MRI-based dataset of cortical thickness. Although the neuroimaging data are cross-sectional, there is overlap between contributing cohorts in their age-distribution. The overlap of the current virtual-histology results with those from our previous longitudinal study ^13^(Vidal-Pineiro et al. 2020) strengthens confidence in these cross-sectional findings. Reliable mapping of gene-expression data to cortical regions was available only from left hemisphere, and derived from a small number of donors. Additionally, detailed gene expression levels mapped to cortical thickness are not currently available for different life-stages.

In summary, we show that neurotypical cortical thinning in early, mid-, and late life is associated with distinct biological pathways related, respectively, to synaptic modeling, neurotransmission, and neurodegeneration. This information provides useful insights into the potential mechanisms underlying age-related changes in cortical organization.

## Acknowledgement

Dr. A Modabbernia is supported by a T32 training grant from the National Institute of Mental Health (MH122394). Dr. D Vidal-Pineiro was by the Norges Forskningsråd (324882). Dr. S Frangou was supported by the National Institutes of Health (R01 MH116147; R01 MH113619; R01 AG058854). Dr. OA Andreasson acknowledges funding from KG Jebsen Stiftelsen, South East Norway Health Authority and Research Council of Norway. Dr. DI Boomsma acknowledges the Royal Netherlands Academy of Science; NWOgrant 480-15-001/674; NWOgrant 051.02.060; NWO/SPI 56-464-14192; NWO 433-09-220. The NeuroIMAGE study was supported by NIH Grant R01MH62873, NWO Large Investment Grant 1750102007010, ZonMW grant 60-60600-97-193, NWO grants 056-13-015 and 433-09-242, and matching grants from Radboud University Nijmegen Medical Center, University Medical Center Groningen and Accare, and Vrije Universiteit Amsterdam. The work was further supported by the EU-AIMs and AIMS-2-TRIALS programmes which receive support from Innovative Medicines Initiative Joint Undertaking Grant No. 115300 and 777394, the resources of which are composed of financial contributions from the European Union’s FP7 and Horizon2020 Programmes, and from the European Federation of Pharmaceutical Industries and Associations (EFPIA) companies’ in-kind contributions, and AUTISM SPEAKS, Autistica and SFARI; and by the Horizon2020 supported programme CANDY Grant No. 847818). PAFIP was supported by the Instituto de Salud Carlos III (00/3095, 01/3129, PI020499, PI14/00639, PI17/01056 and PI14/00918), SENY Fundació Research Grant CI 2005-0308007 and Fundación Marqués de Valdecilla. Instituto de investigación sanitaria Valdecilla (A/02/07, NCT0235832 and NCT02534363). Dr. R Ayesa-Arriola is funded by a Miguel Servet contract from the Carlos III Health Institute (CP18/00003). The BIG database, established in Nijmegen in 2007, is now part of Cognomics, a joint initiative by researchers of the Donders Centre for Cognitive Neuroimaging, the Human Genetics and Cognitive Neuroscience departments of the Radboud University Medical Centre, and the Max Planck Institute for Psycholinguistics. The Cognomics Initiative is supported by the participating departments and centres and by external grants, including grants from the Biobanking and Biomolecular Resources Research Infrastructure (Netherlands) (BBMRI-NL) and the Hersenstichting Nederland. The authors also acknowledge grants supporting their work from the Netherlands Organization for Scientific Research (NWO), i.e. the NWO Brain & Cognition Excellence Program (grant 433-09-229), the Vici Innovation Program (grant 016–130-669 to BF) and #91619115. Additional support is received from the European Community’s Seventh Framework Programme (FP7/2007–2013) under grant agreements n° 602805 (Aggressotype), n° 603016 (MATRICS), n° 602450 (IMAGEMEND), and n° 278948 (TACTICS), and from the European Community’s Horizon 2020 Programme (H2020/2014–2020) under grant agreements n° 643051 (MiND) and n° 667302 (CoCA). Dr. B Franke was also supported by funding from the European Community’s Horizon 2020 Programme (H2020/2014 – 2020) under grant agreements n° 728018 (Eat2beNICE) and n° 847879 (PRIME). They also received relevant funding from the Netherlands Organization for Scientific Research (NWO) for the Dutch National Science Agenda NeurolabNL project (grant 400-17-602). The Study of Health in Pomerania (SHIP) is part of the Community Medicine Research net (CMR) (http://www.medizin.uni-greifswald.de/icm) of the University Medicine Greifswald, which is supported by the German Federal State of Mecklenburg-West Pomerania. MRI scans in SHIP and SHIP-TREND have been supported by a joint grant from Siemens Healthineers, Erlangen, Germany and the Federal State of Mecklenburg-West Pomerania. Dr. H Hulshoff Pol acknowledges BrainSCALE to the Dutch Research Council (NWO 51.02.61, NWO-NIHC Programs of excellence 433-09-220) and UMCU to ZonMW (908.02.123 and 917.46.370) and EU (MRTN-CT-2006-035987). The ENIGMA Working Group acknowledges the NIH Big Data to Knowledge (BD2K) award for foundational support and consortium development (U54 EB020403 to Paul M. Thompson). For a complete list of ENIGMA-related grant support please see here: http://enigma.ini.usc.edu/about-2/funding/. Data collection and sharing for the Pediatric Imaging, Neurocognition and Genetics (PING) Study (National Institutes of Health Grant RC2DA029475) were funded by the National Institute on Drug Abuse and the Eunice Kennedy Shriver National Institute of Child Health & Human Development. A full list of PING investigators is at http://pingstudy.ucsd.edu/investigators.html. The Sydney MAS has been funded by three National Health & Medical Research Council (NHMRC) Program Grants (ID No. ID350833, ID568969, and APP1093083). Dr. H Brodaty also acknowledges funding from the Professor Award (PAH/6635). Indiana 1.5T; Indiana 3T acknowledges NIH grants P30 AG010133, R01 AG019771 and R01 CA129769; Siemens Medical Solutions; the members of the Partnership for Pediatric Epilepsy Research, which includes the American Epilepsy Society, the Epilepsy Foundation, the Epilepsy Therapy Project, Fight Against Childhood Epilepsy and Seizures (F.A.C.E.S.), and Parents Against Childhood Epilepsy (P.A.C.E.); the GE/NFL Head Health Challenge I; the Indiana State Department of Health Spinal Cord and Brain Injury Fund Research Grant Program; a Project Development Team within the ICTSI NIH/NCRR Grant Number RR025761. Dr. A Saykin also acknowledges support from multiple NIH grants (P30 AG010133, P30 AG072976, R01 AG019771, R01 AG057739, U01 AG024904, R01 LM013463, R01 AG068193, T32 AG071444, and U01 AG068057 and U01 AG072177). He has also received support from Avid Radiopharmaceuticals, a subsidiary of Eli Lilly (in kind contribution of PET tracer precursor); Bayer Oncology (Scientific Advisory Board); Eisai (Scientific Advisory Board). QTIM reported National Health and Medical Research Council (NHMRC). Dr. L Nars of BETULA reported a scholar grant from the Knut and Alice Wallenberg (KAW) Foundation. The OATS study has been funded by an NHMRC and Australian Research Council (ARC) Strategic Award Grant of the Ageing Well, Ageing Productively Program (ID No. 401162); NHMRC Project (seed) Grants (ID No. 1024224 and 1025243); NHMRC Project Grants (ID No. 1045325 and 1085606); and NHMRC Program Grants (ID No. 568969 and 1093083). GSP was supported by funding from Siemens Medical Solutions USA, Inc. (Dementia Advisory Board); Springer-Nature Publishing (Editorial Office Support as Editor-in-Chief, Brain Imaging and Behavior). BRCATLAS acknowledges funding from the National Institute for Health Research (NIHR) Maudsley Biomedical Research Centre at South London and Maudsley NHS Foundation Trust and King’s College London. Dr. A Bertolino is a Consultant to HealthLytix, speaker’s honorarium from Lundbeck and Sunovion. Dr. HJ Grabe has received travel grants and speakers honoraria from Fresenius Medical Care, Neuraxpharm, Servier and Janssen Cilag as well as research funding from Fresenius Medical Care. Dr. P Thompson and Dr. N Janahshad received a research grant from Biogen, Inc. for research unrelated to this manuscript.

## References

1 Thambisetty, M. et al. Longitudinal changes in cortical thickness associated with normal aging. Neuroimage 52, 1215–1223, doi:10.1016/j.neuroimage.2010.04.258 (2010).

2 Frangou, S. et al. Cortical thickness across the lifespan: Data from 17,075 healthy individuals aged 3-90 years. Hum Brain Mapp, doi:10.1002/hbm.25364 (2021).

3 Bourgeois, J. P. & Rakic, P. Changes of synaptic density in the primary visual cortex of the macaque monkey from fetal to adult stage. J Neurosci 13, 2801–2820 (1993).

4 Huttenlocher, P. R. Synaptic density in human frontal cortex - developmental changes and effects of aging. Brain Res 163, 195–205, doi:10.1016/0006-8993(79)90349-4 (1979).

5 Huttenlocher, P. R. & Dabholkar, A. S. Regional differences in synaptogenesis in human cerebral cortex. J Comp Neurol 387, 167–178, doi:10.1002/(sici)1096-9861(19971020)387:2<167::aid-cne1>3.0.co;2-z (1997).

6 Yakovlev, P. The myelogenetic cycles of regional maturation of the brain. Regional development of the brain in early life, 3–70 (1967).

7 Grydeland, H. et al. Waves of Maturation and Senescence in Micro-structural MRI Markers of Human Cortical Myelination over the Lifespan. Cereb Cortex 29, 1369–1381, doi:10.1093/cercor/bhy330 (2019).

8 Terry, R. D., DeTeresa, R. & Hansen, L. A. Neocortical cell counts in normal human adult aging. Ann Neurol 21, 530–539, doi:10.1002/ana.410210603 (1987).

9 Marner, L., Nyengaard, J. R., Tang, Y. & Pakkenberg, B. Marked loss of myelinated nerve fibers in the human brain with age. J Comp Neurol 462, 144–152, doi:10.1002/cne.10714 (2003).

10 Morrison, J. H. & Hof, P. R. Life and death of neurons in the aging brain. Science 278, 412–419, doi:10.1126/science.278.5337.412 (1997).

11 French, L. & Paus, T. A FreeSurfer view of the cortical transcriptome generated from the Allen Human Brain Atlas. Front Neurosci 9, 323, doi:10.3389/fnins.2015.00323 (2015).

12 Shin, J. et al. Cell-Specific Gene-Expression Profiles and Cortical Thickness in the Human Brain. Cereb Cortex 28, 3267–3277, doi:10.1093/cercor/bhx197 (2018).

13 Vidal-Pineiro, D. et al. Cellular correlates of cortical thinning throughout the lifespan. Sci Rep 10, 21803, doi:10.1038/s41598-020-78471-3 (2020).

14 Elston, G. N. & Fujita, I. Pyramidal cell development: postnatal spinogenesis, dendritic growth, axon growth, and electrophysiology. Front Neuroanat 8, 78, doi:10.3389/fnana.2014.00078 (2014).

15 Esiri, M. M. Ageing and the brain. J Pathol 211, 181–187, doi:10.1002/path.2089 (2007).

16 Reemst, K., Noctor, S. C., Lucassen, P. J. & Hol, E. M. The Indispensable Roles of Microglia and Astrocytes during Brain Development. Front Hum Neurosci 10, 566, doi:10.3389/fnhum.2016.00566 (2016).

17 Valles, S. L. et al. Function of Glia in Aging and the Brain Diseases. Int J Med Sci 16, 1473–1479, doi:10.7150/ijms.37769 (2019).

18 Thompson, P. M. et al. ENIGMA and global neuroscience: A decade of large-scale studies of the brain in health and disease across more than 40 countries. Transl Psychiatry 10, 100, doi:10.1038/s41398-020-0705-1 (2020).

19 Desikan, R. S. et al. An automated labeling system for subdividing the human cerebral cortex on MRI scans into gyral based regions of interest. Neuroimage 31, 968–980, doi:10.1016/j.neuroimage.2006.01.021 (2006).

20 Wood, S. mgcv: Mixed GAM Computation Vehicle with GCV/AIC/REML smoothness estimation. (2012).

21 Simpson, G. Introducing gratia. From the bottom of the heap (2018).

22 Patel, Y. et al. Virtual Histology of Cortical Thickness and Shared Neurobiology in 6 Psychiatric Disorders. JAMA Psychiatry 78, 47–63, doi:10.1001/jamapsychiatry.2020.2694 (2021).

23 Hawrylycz, M. J. et al. An anatomically comprehensive atlas of the adult human brain transcriptome. Nature 489, 391–399, doi:10.1038/nature11405 (2012).

24 Mancarci, O. & French, L. Homologene: quick access to homologene and gene annotation updates. R package version 1, 68 (2019).

25 Colantuoni, C. et al. Temporal dynamics and genetic control of transcription in the human prefrontal cortex. Nature 478, 519–523, doi:10.1038/nature10524 (2011).

26 Jaffe, A. E. et al. Mapping DNA methylation across development, genotype and schizophrenia in the human frontal cortex. Nat Neurosci 19, 40–47, doi:10.1038/nn.4181 (2016).

27 Trabzuni, D. et al. Quality control parameters on a large dataset of regionally dissected human control brains for whole genome expression studies. J Neurochem 119, 275–282, doi:10.1111/j.1471-4159.2011.07432.x (2011).

28 Consortium, G. T. Human genomics. The Genotype-Tissue Expression (GTEx) pilot analysis: multitissue gene regulation in humans. Science 348, 648–660, doi:10.1126/science.1262110 (2015).

29 Sliz, E. et al. A variant near DHCR24 associates with microstructural properties of white matter and peripheral lipid metabolism in adolescents. Mol Psychiatry 26, 3795–3805, doi:10.1038/s41380-019-0640-9 (2021).

30 Yu, G., Wang, L. G., Han, Y. & He, Q. Y. clusterProfiler: an R package for comparing biological themes among gene clusters. OMICS 16, 284–287, doi:10.1089/omi.2011.0118 (2012).

31 Whitaker, K. J. et al. Adolescence is associated with genomically patterned consolidation of the hubs of the human brain connectome. Proc Natl Acad Sci U S A 113, 9105–9110, doi:10.1073/pnas.1601745113 (2016).

32 Golbs, A., Nimmervoll, B., Sun, J. J., Sava, I. E. & Luhmann, H. J. Control of programmed cell death by distinct electrical activity patterns. Cereb Cortex 21, 1192–1202, doi:10.1093/cercor/bhq200 (2011).

33 Parker, N. et al. Corticosteroids and Regional Variations in Thickness of the Human Cerebral Cortex across the Lifespan. Cereb Cortex 30, 575–586, doi:10.1093/cercor/bhz108 (2020).

34 Rojas, A. M., Fuentes, G., Rausell, A. & Valencia, A. The Ras protein superfamily: evolutionary tree and role of conserved amino acids. J Cell Biol 196, 189–201, doi:10.1083/jcb.201103008 (2012).

35 Yamamoto, A. & Behl, C. Human Nck-associated protein 1 and its binding protein affect the metabolism of beta-amyloid precursor protein with Swedish mutation. Neurosci Lett 316, 50–54, doi:10.1016/s0304-3940(01)02370-9 (2001).

36 Liu, F. F., Zhang, Z., Chen, W., Gu, H. Y. & Yan, Q. J. Regulatory mechanism of microRNA-377 on CDH13 expression in the cell model of Alzheimer’s disease. Eur Rev Med Pharmacol Sci 22, 2801–2808, doi:10.26355/eurrev_201805_14979 (2018).

37 Zhang, X., Huang, T. Y., Yancey, J., Luo, H. & Zhang, Y. W. Role of Rab GTPases in Alzheimer’s Disease. ACS Chem Neurosci 10, 828–838, doi:10.1021/acschemneuro.8b00387 (2019).

38 Vogrinc, D., Goricar, K., Kunej, T. & Dolzan, V. Systematic Search for Novel Circulating Biomarkers Associated with Extracellular Vesicles in Alzheimer’s Disease: Combining Literature Screening and Database Mining Approaches. J Pers Med 11, doi:10.3390/jpm11100946 (2021).

39 Olah, M. et al. A transcriptomic atlas of aged human microglia. Nat Commun 9, 539, doi:10.1038/s41467-018-02926-5 (2018).

40 Wong, A. P. et al. Inter-Regional Variations in Gene Expression and Age-Related Cortical Thinning in the Adolescent Brain. Cereb Cortex 28, 1272–1281, doi:10.1093/cercor/bhx040 (2018).

