## Supplemental for "Dissociable Cellular and Genetic Mechanisms of Cortical Thinning at Different Life Stages"

**Supplementary Data**

A. **Supplementary Methods**
1. **Neuroimaging Dataset**

Supplementary Table 1. Demographic Characteristics of the Contributing Cohorts

Supplementary Figure 1. Age-Distribution in the Contributing Cohorts

Supplementary Table 2. Screening Process and Eligibility Criteria, Scanner, Image Acquisition Parameters and Image Segmentation Software

2. **Quality Assurance for the Neuroimaging Data**
3. **Sensitivity Analyses of the General Additive Mixed Models**

Supplementary Table 3. Correlation Between the Cortical Thinning Profiles from the Main Analysis and Cortical Thinning Profile Obtained Using Different Number of Knots of the Generalized Additive Mixed Models

4. **Cell-Specific Gene Expression Profiling**
5. **Study Specific Gene Expression Database and Gene Co-expression Networks**
6. **Permutation Analyses for the Correlation Between Lifespan Thinning and Cell-Specific Gene Expression Profiles**
7. **Code Availability**
B. **Supplementary Results**
1. **Cortical Thinning Estimates Across the Lifespan**

Supplementary Figure 2. Estimates of Cortical Thinning Across the Lifespan

2. **Sensitivity Analyses for the Number of Knots in the General Additive Mixed Models for Cortical Thinning on Virtual Histology**

Supplementary Figure 3. Effect of Varying Number of Knots in the Generalized Additive Mixed Models on Virtual Histology

3. **Association of life-stage inter-cortical thinning profiles with gene co-expression and enrichment**

Supplementary Figure 4. Cnetplot for the Linkages of Genes and Biological Processes in Early-Life

Supplementary Figure 5. Cnetplot for the Linkages of Genes and Cellular Components in Early-Life

Supplementary Figure 6. Cnetplot for the Linkages of Genes and Biological Processes in Mid-Life

Supplementary Figure 7. Cnetplot for the Linkages of Genes and Cellular Components in Mid-Life

Supplementary Figure 8. Cnetplot for the Linkages of Genes and Biological Processes in Late-Life

Supplementary Figure 9. Cnetplot for the Linkages of Genes and Cellular Components in Late-Life

4. **Sensitivity Analyses for Gene Ontology**
C. **Supplementary References**
A. **Supplementary Methods**
1. **Neuroimaging Dataset**

| **Supplementary Table 1. Demographic Characteristics of the Included Cohorts** | | | | |
| --- | --- | --- | --- | --- |
| **Cohort Name** | **Sample Size** | **Number of Males** | **Number of Females** | **Age**  **Mean (SD)** |
| **Betula** | 231 | 105 | 126 | 62.5 (12.4) |
| **BIG - 1.5T** | 1319 | 657 | 662 | 28.3 (14.3) |
| **BIG -  3T** | 1125 | 493 | 632 | 24.0 (8.3) |
| **BIL&GIN** | 452 | 220 | 232 | 26.7 (7.7) |
| **Bonn** | 175 | 175 | 0 | 38.8 (6.5) |
| **BRAINSCALE** | 172 | 102 | 70 | 10.1 (1.4) |
| **BRCATLAS** | 163 | 84 | 79 | 39.7 (17.2) |
| **CAMH** | 141 | 72 | 69 | 43.6 (19.3) |
| **Cardiff** | 265 | 78 | 187 | 25.6 (7.8) |
| **CLiNG** | 323 | 132 | 191 | 25.2 (5.3) |
| **FIDMAG** | 123 | 54 | 69 | 37.5 (10.1) |
| **GSP** | 1924 | 854 | 1070 | 26.8 (16.4) |
| **HUBIN** | 102 | 69 | 33 | 42.0 (8.8) |
| **IDIVAL 3** | 104 | 63 | 41 | 30.2 (7.8) |
| **IMAGEN** | 1722 | 854 | 868 | 14.5 (0.4) |
| **Indiana 3T** | 184 | 83 | 101 | 28.1 (20.3) |
| **Leiden** | 572 | 279 | 293 | 16.9 (4.8) |
| **MAS** | 385 | 176 | 209 | 78.5 (4.7) |
| **Muenster** | 744 | 323 | 421 | 35.2 (12.1) |
| **NCNG** | 345 | 110 | 235 | 51.4 (16.9) |
| **NeuroIMAGE** | 252 | 115 | 137 | 16.7 (3.4) |
| **Neuroventure** | 137 | 62 | 75 | 13.7 (0.6) |
| **NTR 2** | 112 | 42 | 70 | 33.8 (10.4) |
| **OATS 3** | 116 | 64 | 52 | 69.5 (4.0) |
| **Olin** | 582 | 231 | 351 | 35.9 (13.0) |
| **PING** | 109 | 53 | 56 | 9.0 (3.5) |
| **QTIM** | 308 | 96 | 212 | 22.6 (3.3) |
| **SHIP 2** | 306 | 172 | 134 | 54.5 (12.3) |
| **SHIP TREND** | 628 | 355 | 273 | 49.9 (13.7) |
| **TOP** | 303 | 159 | 144 | 35.4 (9.9) |
| **Sydney** | 157 | 65 | 92 | 39.1 (22.1) |
| **UMCU 1 (CTR) 3T** | 144 | 69 | 75 | 43.9 (14.0) |
| **UMCU 2 (1.5T)** | 278 | 158 | 120 | 32.9 (12.5) |
| **UNIBA** | 130 | 67 | 63 | 27.4 (9.1) |
| **UPENN** | 115 | 42 | 73 | 36.8 (13.1) |
| **Abbreviations of studies**: Betula = Swedish longitudinal study on aging, memory, and dementia; BIG = Brain Imaging Genetics; BIL&GIN = a multimodal multidimensional database for investigating hemispheric specialization; Bonn = University of Bonn; BrainSCALE=Brain Structure and Cognition: an Adolescence Longitudinal twin study; BRCATLAS = NIHR Biomedical Research Centre/ Mapping the relationship between the white matter and executive function across the adult lifespan; CAMH = Centre for Addiction and Mental Health; Cardiff = Cardiff University; CLiNG = Clinical Neuroscience Göttingen; FIDMAG = Fundación para la Investigación y Docencia Maria Angustias Giménez; GSP = Brain Genomics Superstruct Project; HUBIN = Human Brain Informatics; IDIVAL = Valdecilla Biomedical Research Institute; IMAGEN = the IMAGEN Consortium; Indiana = Indiana University School of Medicine; Leiden = Leiden University; MAS = Memory and Ageing Study; Muenster = Muenster University; NCNG = Norwegian Cognitive NeuroGenetics sample; NeuroIMAGE = Dutch part of the International Multicenter ADHD Genetics (IMAGE) study; Neuroventure: the imaging part of the Co-Venture Trial funded by the Canadian Institutes of Health Research (CIHR); NTR = Netherlands Twin Register; OATS = Older Australian Twins Study; Olin = Olin Neuropsychiatric Research Center; PING = Pediatric Imaging, Neurocognition, and Genetics; QTIM = Queensland Twin Imaging; SHIP-2 and SHIP TREND = Study of Health in Pomerania; Sydney = University of Sydney; TOP = Tematisk Område Psykoser (Thematically Organized Psychosis Research); UMCU = Universitair Medisch Centrum Utrecht; UNIBA = University of Bari Aldo Moro; UPENN=University of Pennsylvania | | | | |

**Supplementary Figure 1. Age-distribution of the Contributing Cohorts**

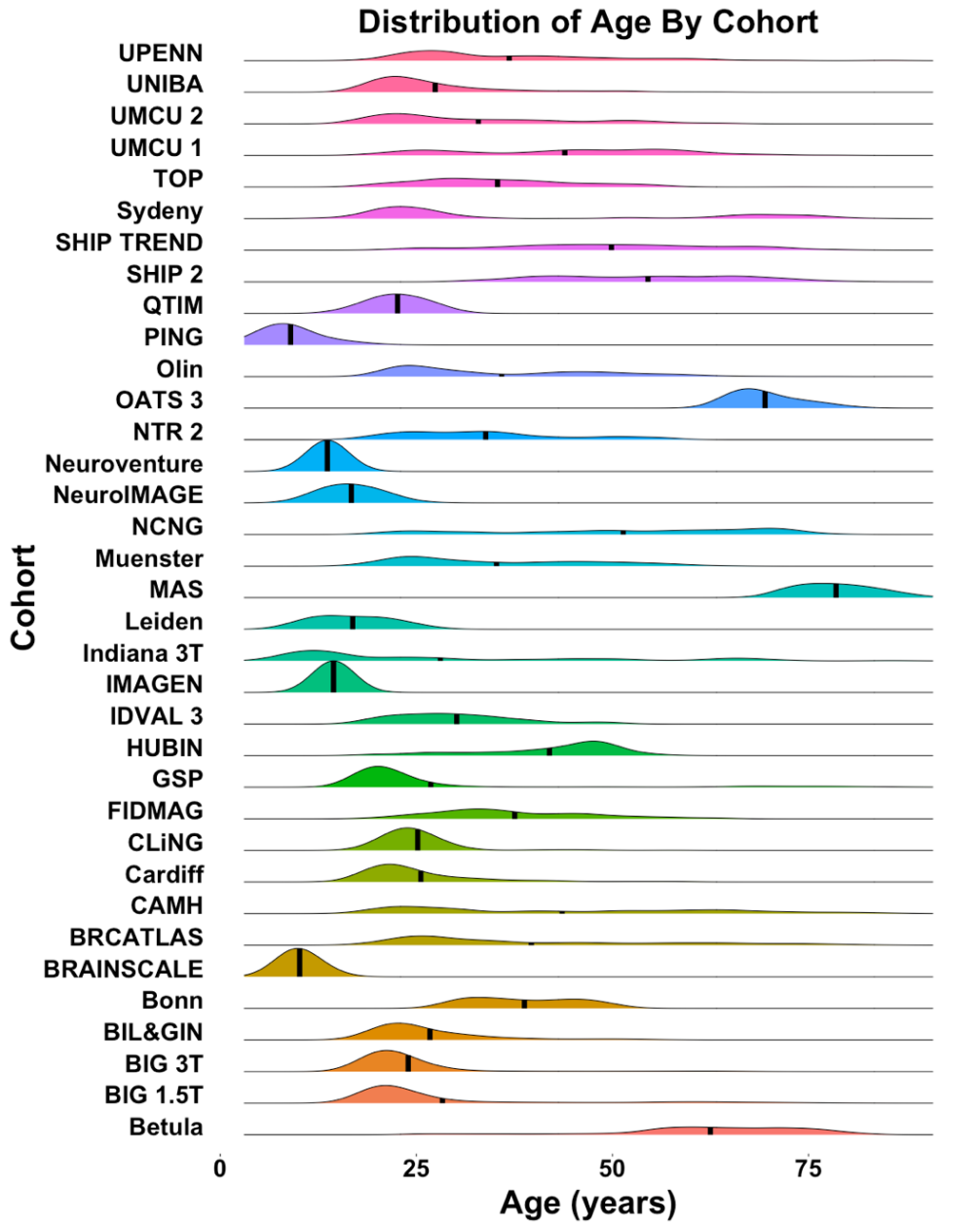

| **Supplementary Table 2. Screening Process and Eligibility Criteria, Scanner, Image Acquisition Parameters and Image Segmentation Software** | | | | | |
| --- | --- | --- | --- | --- | --- |
| **Sample** | **Screening Process** | **Eligibility Criteria** | **Magnet strength/ Scanner Vendor** | **Acquisition parameters** | **Freesurfer version** |
| **Betula** | Personal Interview | No head trauma, no medical, neurological or psychiatric history, no lifetime alcohol or substance abuse, no previous or current use of psychotropic medication. | 3T General Electric Discovery  MR750 | T1-weighted MPRAGE; TR/TE/TI/FA=8.1240 ms/3.2000 ms/450ms/ 12°; image matrix = 256x256 | 5.3 |
| **BIG** | Questionnaire for psychiatric history | No history of somatic disease potentially affecting the brain, current or past psychiatric or neurological disorder, medication (except hormonal contraceptives) or illicit drug use during the past 6 months, history of substance abuse, current or past alcohol dependence, pregnancy, lactation, menopause, or MRI contraindications. | 1,5 T Siemens Sonata and Avanto and 3 T Siemens Trio, TimTrio and Skyra | T1-weighted 3D MPRAGE; TR/TE/TI/sagittal slices = 1940-2730 ms/850-110 ms/2.92-4.58 ms; 176-192 sagittal slices; voxel size= 1.0x1.0x1.0 mm^3^ | 5.3 |
| **BIL&GIN** | Personal Interview | No head trauma, no current neurological or psychiatric disorders, no current use of psychotropic medication, IQ>70. | 3T Phillips ACHIEVA | T1 - weighted 3D; TR/TE/TI/FA=20 ms/4.6 ms/800ms/ 10°; turbo field echo factor = 65; sense factor = 2; matrix size = 256x256x180mm 3 ; voxel size= 1.0x1.0x1.0 mm^3^ | 5.3 |
| **Bonn** | Personal interview | No head trauma, no medical, neurological or psychiatric history, no previous or current use of psychotropic medication. | 3T Siemens Trio | TR/TE/FA= 1570-1660ms/2.75-3.42ms/8-9° | 5.3 |
| **BrainSCALE** | Personal interview | No head trauma, no medical, neurological or psychiatric history, no lifetime alcohol or substance abuse, no previous or current use of psychotropic medication. | 1.5T Philips Achieva | T1-weighted 3D SPGR; TR/TE/ FA= 30 ms/4.6 ms/30°; image matrix=256x256; 160**–**180 contiguous coronal slices; voxel size=1 x 1 x 1.2 mm^3^ | 5.1 |
| **BRCATLAS** | Telephone interview | No head trauma, no medical, neurological or psychiatric history, no lifetime alcohol or substance abuse, no mild cognitive impairment, no previous or current use of psychotropic medication, IQ>75. | 3T GE Signa | T1-weighted 3D; TR/TE/TI/FA= 6.9 ms/2.8 ms/650 ms/8°; Image matrix = 256 x 256 x 180mm^3^; voxel size=1mm^3^ | 5.3 |
| **CAMH** | SCID | No head trauma, no neurological or psychiatric history, no alcohol or substance abuse preceding 6 months, no previous or current use of psychotropic medication, IQ>75. No history of any psychotic disorders in 1st degree family members. | 1.5 T GE (echospeed) | 124 Axial inversion recovery–prepared spoiled gradient recall images, 1.5-mm-thick slice acquisition TE/TR/TI/FA=5.3ms/12.3ms/300.0ms/20°. | 5.3 |
| **Cardiff** | MINI | No head trauma, no medical history, including neurological and psychiatric history, no alcohol or substance abuse in the preceding 6 months, no previous or current use of psychotropic medication. | 3T General Electric Signa | T1-weighted 3D FSPGR; TR/TE/TI/FA=7.9ms/3.0ms/450ms/20^o^;  Image matrix= 256 × 192 x 172; voxel size=1mm^3^ | 5.3 |
| **CLiNG** | Personal Interview | No head trauma, no medical, neurological or psychiatric history, no lifetime alcohol or substance abuse, no previous or current use of psychotropic medication, IQ>75. No history of any psychiatric disorders in 1st degree family members. | 3T Siemens Tim Trio | T1-weighted 3D MPRAGE; TR/TE/TI/FA=2250 ms/3.26 ms/900 ms/9°; image matrix = 256 x 256; 192 sagittal slices; voxel size= 1 mm^3^ | 5.3 |
| **FIDMAG** | Personal interview; structured interview in part of the sample | No head trauma, no medical, neurological or psychiatric history, no lifetime alcohol or substance abuse, no previous or current use of psychotropic medication, IQ>70. | 1.5 T General Electric Signa | T1-weighted MPRAGE; TR/TE/FA=2000 ms/4 ms/ 9^o^; image matrix=512 x 512; 180 contiguous sagittal slices; voxel size=0.56 x 0.56 x 1 mm3 | 5.3 |
| **GSP** | Structured phone screen and study specific self-report battery and clinical screen | No head trauma, no medical, neurological or psychiatric history, no lifetime alcohol or substance abuse, no current use of psychotropic medication, normal brain anatomy following brain scan. | 3T Siemens Tim Trio | T1-weighted 3D multi-echo MPRAGE; TR/TE/TI/FA =2200 ms/1.54-7 ms/  1100/7 ^o^; voxel size=1.2x1.2x1.2 mm | 5.3 |
| **HUBIN** | SCID-I | No head trauma, no medical, neurological or psychiatric history, no lifetime alcohol or substance abuse, no previous or current use of psychotropic medication, IQ>75. No history of any psychiatric disorders in 1st degree family members. | 1.5 T General Electric Signa | T1-weighted SPGR; TR/TE/FA= 24 ms/6 ms/35 ^o^; 124 coronal slices; voxel size 0.86 x 0.86 x 1.50 mm3. | 5.3 |
| **IDIVAL (3)** | Personal Interview | No lifetime history of Axis I psychiatric disorders, no mild cognitive | 3T Phillips Achieva | T1-weighted SPGR; TR/TE/FA=3000 ms/4.6 ms/8^o^; image matrix=321x312; voxel size=1mm^3^; sagittal plane acquisition | 5.3 |
| **IMAGEN** | DAWBA questionnaire clinician interview | No head trauma, no medical, neurological or psychiatric history, no previous or current use of psychotropic medication. IQ>75 | 3T Siemens Verio and TimTrio, Philips Achieva, General Electric Signa Excite, and Signa HDx | T1-weighted 3D MPRAGE; TR/TE/TI/FA=2300ms/3030ms/900ms/9^o^;  Image matrix= 256 × 256;192 sagittal slices; voxel size=1mm^3^ | 5.3 |
| **Indiana 3T** | Personal interview  Structured phone screen | No head trauma, no medical, neurological or psychiatric history, no alcohol or substance abuse in the preceding 6 months, no previous or current use of psychotropic medication, IQ>75. | 3T Siemens Skyra | T1-weighted MPRAGE; TR/TE/FA=2300 ms/2.95 ms/ 9^o^; image matrix=256  x 240; 176 contiguous sagittal slices | 5.1 |
| **Leiden** | Self-report | No psychiatric or neurological disorders, no use of psychotropic medications | 3T Philips Achieva | T1-weighted 3D SPGR; TR/TE = 9.76 ms/4.59 ms; image matrix=256x256; 160**–**180 contiguous coronal slices; voxel size=0.875x 0.875 x 1.2 mm^3^ | 5.3 |
| **MAS** | Personal interview | No head trauma, no diagnosis of dementia, schizophrenia, bipolar disorder no psychotic symptoms, no neurological disorder, no mild cognitive impairment, or intellectual disability. | 3T Philips Achieva Quasar Dual | TR/TE = 6.39 ms/2.9 ms; 190 coronal slices; voxel size = 1mm^3^ | 5.3 |
| **Muenster** | SCID | No head trauma, no medical, neurological or psychiatric history, no lifetime alcohol or substance abuse, no previous or current use of psychotropic medication, IQ>75. | 3T Phillips Intera | T1 weighted TFE: TR/TE/FA= 7.4 ms/3.4 ms/9°; image matrix = 256x204x160mm^3^; voxel size=0.5mm^3^; sagittal plane acquisition | 5.3 |
| **NCNG** | Personal interview | No head trauma, no medical, neurological or psychiatric history, no lifetime alcohol or substance abuse, no mild cognitive impairment, no previous or current use of  psychotropic medication, IQ>84. | 1.5T Siemens Avanto  1.5T Siemens Sonata | T1-weighted 3D MPRAGE; TR/TE/TI/FA= 2400 ms/3.61 ms/1000 ms/8°; image matrix=192x192; 160 sagittal slices; voxel size=1.25 mm^3^  T1-weighted 3D MPRAGE; TR/TE/TI/FA= 2730 ms/3.43 ms/1000 ms/7°; image matrix=256x256; 128 sagittal slices; voxel size=1 mm^3^ | 5.3 |
| **NeuroIMAGE** | KSADS-PL | No head trauma, no mild cognitive impairment, neurological or psychiatric history, no previous or current use of psychotropic medication, IQ>75. No history of any psychiatric disorders in 1st and 2nd degree family members. | 1.5 T Siemens AVANTO (Donders Centre for Cognitive Neuroimaging)  1.5 T Siemens SONATA (VU University Amsterdam) | MPRAGE 176 sagittal slices, repetition time=2,730ms, echo time=2.95ms, voxel size=1.0x1.0x1.0mm, field of view=256 mm | 5.3 |
| **Neuroventure** | DAWBA and BSI | No head trauma, no medical, neurological or psychiatric history, no lifetime alcohol or substance abuse, no previous or current use of psychotropic medication, IQ>75. | 3T SIEMENS TrioTim | T1-weighted 3D MPRAGE; TR/TE/ FA= 2300 ms/2.96 ms/9^o^; image matrix= 256x256; voxel size= 1.0x1.0x1.0 mm^3^ | 5.3 |
| **NTR (2)** | MINI, BDI, STAI, STAS, YBOCS | No head trauma, no previous or current use of psychotropic medication, normal IQ. | 3T Philips Intera | T1-weighted 3D MPRAGE; TR/TE/FA=9.64 ms/4.60 ms/8 ^o^; image matrix=256 x 256; 182 coronal slices; voxel size=1 x1x1.2 mm^3^ | 5.1 |
| **OATS (1-4)** | Personal interview | No head trauma, no current diagnosis of a psychotic disorder, no neurological disorder, no malignancy (other than skin cancer) or other severe medical comorbidity, no mild cognitive impairment, or intellectual disability. | 1.5T Philips Gyroscan, Siemens Magnetom Avanto, Siemens Sonata; 3T Philips Achieva Quasar Dual, a | T1-weighted 3D acquisition; TR/TE/TI/FA=15370 ms/3.24 ms/780 ms/8^o^; 144 slices; voxel size=1 x 1 x 1.5 mm^3^ | 5.3 |
| **OLIN** | SCID I | No head trauma, no medical, neurological or psychiatric history, no alcohol or substance abuse preceding 6 months, never prescribed with psychotropic medication, IQ>75. | 3T Siemens Alegra | T1-weighted 3D MPRAGE; TR/TE/TI/FA= 2300 ms/2.91 ms/900 ms/9^o^; image matrix= 256x240x192; 160 sagittal slices; voxel size= 1.0x1.0x1.2 mm^3^ | 5.1 |
| **PING** | Personal interview | No lifetime history of major developmental, psychiatric, or neurological disorders, brain injury, or other medical conditions that affect development. Individuals born  earlier than 36 weeks of gestational age were excluded. | 3T Philips Achieva  3T GE SIGNA  3T Siemens TrioTim  3T Siemens TrioTim  3T General Electric Discovery  MR750 | T1-weighted 3D IR-GRE; TR/TE/TI/FA= 8.1 ms/3.5 ms/640 ms/9^o^ | 5.3 |
| **QTIM** | CIDI | No head trauma, no medical history, neurological and psychiatric history, no alcohol or substance abuse in the preceding 6 months, no antidepressant medication or medication affecting cognition. | 4T Bruckner | T1-weighted 3D MPRAGE: TR/TE/TI/FA = 1500 ms/3.35 ms/ 700 ms/ 8°; image matrix= 256 × 256 × 256 or 256 × 256 × 240; 256 coronal slices; voxel size= 0.9 mm^3^ | 5.1 |
| **SHIP-2** | Personal Interview | No head trauma, no neurological and psychiatric history, no risky alcohol consumption (cut-offs: males: >= 60g alcohol per day, females: >=30 g alcohol per day) in the preceding 30 days, no current use of psychotropic medication. Exclusion of school leavers without degree. Exclusion of strong MRI artifacts and inhomogeneities | 1.5T Siemens Avanto | T1-weighted 3D MPRAGE; TR/TE/ FA=1900ms/3.4ms/15^o^; voxel size=1mm^3^ | 5.3 |
| **SHIP-TREND** | Personal Interview | No head trauma, no neurological and psychiatric history, no risky alcohol consumption (cut-offs: males: >= 60g alcohol per day, females: >=30 g alcohol per day) in the preceding 30 days, no current use of psychotropic medication. Exclusion of school leavers without degree. Exclusion of strong MRI artifacts and inhomogeneities | 1.5T Siemens Avanto | T1-weighted 3D MPRAGE; TR/TE/ FA=1900ms/3.4ms/15^o^; voxel size=1 mm^3^ | 5.3 |
| **Sydney** | SCID | No head trauma, no medical history, neurological or psychiatric history, no alcohol or substance abuse preceding 6 months, as well as no previous or current use of psychotropic medication, IQ>75. | 3T General Electric Discovery MR750 | T1-weighted 3D MPRAGE; TR/TE/FA= 7264ms/ 2784ms/15^o^; image matrix  =256 x 256 x 196; voxel size=0.9mm^3^ | 5.1 |
| **TOP** | PRIME-MD | No head trauma, no organic or other psychotic disorder (ICD codes 290-299), no substance abuse in the preceding 6 months, no previous or current use of psychotropic medication, IQ>75. No history of any psychiatric disorders in 1st degree family members. | 1.5T Siemens Magnetom Sonata | T1-weighted SPGR; TR/TE/TI/FA=2730 ms/3.93 ms/1000 ms/71^o^; voxel size = 1.33x0.94x1mm^3^; sagittal plane acquisition | 5.3 |
| **UMCU** | CASH | No head trauma, no medical, neurological or psychiatric history, no lifetime alcohol or substance abuse, no previous or current use of psychotropic medication, IQ>75. No history of any psychiatric disorders in 1st degree family members. | 1.5T Philips Intera and Achieva | T1-weighted 3D FFE; TE/TR/FA= 4.6 ms/0 ms**/ 0˚;** 160-180 contiguous coronal slices; voxel size=1x1x1.2 mm^3^ | 5.1 |
| **UNIBA** | SCID-NP | No head trauma, no medical, neurological or psychiatric history, no lifetime alcohol or substance abuse, no previous or current use of psychotropic medication, IQ>75. No history of any psychiatric disorders in 1st degree family members. | 3T General Electric | T1-weighted 3D SPGR; TE/FA = min full/ 6°; image matrix= 256×256 x124 | 5.3 |
| **UPENN** | SCID | No head trauma, no medical history, including neurological and psychiatric history, no alcohol or substance abuse preceding 6 months, no previous or current use of psychotropic medication, IQ>75. No history of any psychiatric disorders in 1st degree family members. | 3T Siemens Tim Trio | T1-weighted 3D MPRAGE; TR/TE/TI/FA=1810 ms/3.51 ms/1100 ms/9^o^; image matrix= 256 × 192;160 axial slices | 5.3 |
| Abbreviations of Terms: BDI = Behavioural Descriptive Interview; BSI = Brief Symptom Inventory; CASH = Comprehensive assessment of symptoms and history; CDR = Clinical Dementia Rating; CIDI = Composite International Diagnostic Interview; DAWBA = Development and Well-Being Assessment; DISC-IV = Diagnostic Interview Schedule for Children; DSM = Diagnostic and Statistical Manual of Mental Disorders (DSM); FA=flip angle; FSPGR=fast spoiled gradient echo sequence; GRE=spoiled gradient echo sequence; ICD= International Classification of Diseases; IR= inversion recovery; KSADS-PL= Kiddie Schedule for Affective Disorders and Schizophrenia-Present and Lifetime; MADRS = Montgomery-Asberg Depression Rating Scale; MINI = Mini International Neuropsychiatric Interview; MMSE = Mini Mental State Exam; PRIME-MD = Primary Care Evaluation of Mental Disorder; SCID = Structured Clinical Interview for DSM Disorders; SCID-I/NP = SCID Non-Patient version; SPGR=spoiled gradient recalled sequence; STAI = State-Trait Anxiety Inventory; STAS = State-trait anger scale; TE=echo time; TI=inversion time; TR=repetition time; TFE=turbo field echo sequence; YBOCS = Yale-Brown Obsessive Compulsive Scale  **Abbreviations of studies**: Betula = Swedish longitudinal study on aging, memory, and dementia; BIG = Brain Imaging Genetics; BIL&GIN = a multimodal multidimensional database for investigating hemispheric specialization; Bonn = University of Bonn; BrainSCALE=Brain Structure and Cognition: an Adolescence Longitudinal twin study; BRCATLAS = NIHR Biomedical Research Centre/ Mapping the relationship between the white matter and executive function across the adult lifespan; CAMH = Centre for Addiction and Mental Health; Cardiff = Cardiff University; CLiNG = Clinical Neuroscience Göttingen; FIDMAG = Fundación para la Investigación y Docencia Maria Angustias Giménez; GSP = Brain Genomics Superstruct Project; HUBIN = Human Brain Informatics; IDIVAL = Valdecilla Biomedical Research Institute; IMAGEN = the IMAGEN Consortium; Indiana = Indiana University School of Medicine; Leiden = Leiden University; MAS = Memory and Ageing Study; Muenster = Muenster Neuroimaging Cohort; NCNG = Norwegian Cognitive NeuroGenetics sample; NeuroIMAGE = Dutch part of the International Multicenter ADHD Genetics (IMAGE) study; Neuroventure: the imaging part of the Co-Venture Trial funded by the Canadian Institutes of Health Research (CIHR); NTR = Netherlands Twin Register; OATS = Older Australian Twins Study; Olin = Olin Neuropsychiatric Research Center; PING = Pediatric Imaging, Neurocognition, and Genetics; QTIM = Queensland Twin Imaging; SHIP-2 and SHIP TREND = Study of Health in Pomerania; Sydney = University of Sydney; TOP = Tematisk Område Psykoser (Thematically Organized Psychosis Research); UMCU = Universitair Medisch Centrum Utrecht; UNIBA = University of Bari Aldo Moro; UPENN=University of Pennsylvania | | | | | |

**2.	Quality Assurance for the Neuroimaging Data**

Site analysts visually inspected the scans to remove those with radiological findings. Then the quality of the scans was rated using a visual grading system. Scans considered “unacceptable” were excluded at the level of each individual site. Scans rated as “good” or “fair” were subjected to parcellation. Following parcellation, site analysts inspected the 34 bilateral cortical Desikan-Killiany atlas segmentations for each participant. Visual inspection was conducted to assess the success of the extraction of the cortical grey matter ribbon, to identify regional boundary errors on the cortical surface, and ensure the accuracy of anatomical labels. Images were inspected slice by slice using orthogonal and external surface displays. Following qualitative assessment, the regional parcellations were marked on a binary “pass” or “failed” scale. Participants with regions marked as “failed” were removed. After these two steps, each site forwarded data from individuals with successful parcellations to the Icahn School of Medicine at Mount Sinai, where regional cortical thickness estimates identified as outliers using five median absolute deviations (MAD) were removed.

3. **Sensitivity Analyses of the General Additive Mixed Models**

Correlations between cortical thinning measures extracted with different number of knots ranged 0.75-0.99 (mean r=0.96) (Supplementary Table 3) demonstrating the robustness of the results to the GAMM parameters**.**

| **Supplementary Table 3. Correlation Between the Cortical Thinning Profile from the Main Analysis and Cortical Thinning Profile Obtained from Changing the Number of Knots of the Generalized Additive Mixed Models** | | | | | | | | | |
| --- | --- | --- | --- | --- | --- | --- | --- | --- | --- |
| **Age, Years** | **Number of Knots** | | | | | | | | |
|  | **6** | **7** | **8** | **9** | **10** | **12** | **15** | **20** | **40** |
| 4 | 0.77 | 0.76 | 0.84 | 0.95 | 0.91 | 0.88 | 0.92 | 0.92 | 0.93 |
| 5 | 0.85 | 0.83 | 0.89 | 0.97 | 0.94 | 0.92 | 0.95 | 0.95 | 0.96 |
| 6 | 0.91 | 0.89 | 0.93 | 0.98 | 0.96 | 0.95 | 0.96 | 0.97 | 0.97 |
| 7 | 0.95 | 0.93 | 0.96 | 0.98 | 0.97 | 0.97 | 0.96 | 0.97 | 0.97 |
| 8 | 0.97 | 0.95 | 0.97 | 0.99 | 0.98 | 0.98 | 0.96 | 0.97 | 0.97 |
| 9 | 0.98 | 0.97 | 0.98 | 0.99 | 0.99 | 0.98 | 0.97 | 0.97 | 0.97 |
| 10 | 0.99 | 0.98 | 0.99 | 0.99 | 0.99 | 0.98 | 0.98 | 0.98 | 0.98 |
| 11 | 0.99 | 0.98 | 0.99 | 0.99 | 0.99 | 0.98 | 0.99 | 0.98 | 0.98 |
| 12 | 0.99 | 0.99 | 0.99 | 0.99 | 0.99 | 0.99 | 0.99 | 0.98 | 0.98 |
| 13 | 0.99 | 0.99 | 0.99 | 0.99 | 0.99 | 0.99 | 0.98 | 0.99 | 0.98 |
| 14 | 0.99 | 0.99 | 0.99 | 0.99 | 0.99 | 0.99 | 0.98 | 0.98 | 0.98 |
| 15 | 0.99 | 0.99 | 0.99 | 0.99 | 0.99 | 0.99 | 0.98 | 0.98 | 0.98 |
| 16 | 0.99 | 0.99 | 0.99 | 0.99 | 0.99 | 0.99 | 0.98 | 0.97 | 0.97 |
| 17 | 0.99 | 0.99 | 0.99 | 0.99 | 0.99 | 0.99 | 0.98 | 0.97 | 0.97 |
| 18 | 0.99 | 0.99 | 1 | 0.99 | 1 | 0.99 | 0.99 | 0.98 | 0.98 |
| 19 | 0.99 | 1 | 0.99 | 0.99 | 1 | 0.99 | 1 | 0.99 | 0.99 |
| 20 | 0.99 | 1 | 0.99 | 0.99 | 1 | 0.99 | 0.99 | 0.99 | 0.98 |
| 21 | 0.99 | 1 | 0.99 | 0.99 | 0.99 | 0.99 | 0.98 | 0.97 | 0.97 |
| 22 | 0.99 | 0.99 | 0.98 | 0.99 | 0.99 | 0.99 | 0.97 | 0.96 | 0.96 |
| 23 | 0.99 | 0.99 | 0.98 | 0.99 | 0.98 | 0.99 | 0.96 | 0.96 | 0.96 |
| 24 | 0.99 | 0.98 | 0.98 | 0.99 | 0.98 | 0.99 | 0.97 | 0.97 | 0.96 |
| 25 | 0.99 | 0.97 | 0.98 | 0.98 | 0.98 | 0.97 | 0.97 | 0.96 | 0.95 |
| 26 | 0.99 | 0.96 | 0.98 | 0.98 | 0.98 | 0.96 | 0.96 | 0.94 | 0.94 |
| 27 | 0.99 | 0.95 | 0.99 | 0.98 | 0.98 | 0.95 | 0.95 | 0.94 | 0.94 |
| 28 | 0.98 | 0.94 | 0.98 | 0.97 | 0.99 | 0.94 | 0.94 | 0.95 | 0.95 |
| 29 | 0.98 | 0.95 | 0.98 | 0.97 | 0.98 | 0.94 | 0.94 | 0.96 | 0.96 |
| 30 | 0.97 | 0.95 | 0.97 | 0.97 | 0.98 | 0.95 | 0.95 | 0.96 | 0.95 |
| 31 | 0.96 | 0.96 | 0.95 | 0.97 | 0.97 | 0.96 | 0.97 | 0.95 | 0.94 |
| 32 | 0.96 | 0.97 | 0.94 | 0.97 | 0.96 | 0.97 | 0.96 | 0.93 | 0.93 |
| 33 | 0.96 | 0.98 | 0.92 | 0.98 | 0.95 | 0.96 | 0.91 | 0.91 | 0.91 |
| 34 | 0.95 | 0.98 | 0.91 | 0.98 | 0.94 | 0.94 | 0.87 | 0.87 | 0.88 |
| 35 | 0.95 | 0.97 | 0.9 | 0.98 | 0.94 | 0.91 | 0.83 | 0.84 | 0.86 |
| 36 | 0.94 | 0.95 | 0.9 | 0.98 | 0.93 | 0.9 | 0.83 | 0.83 | 0.84 |
| 37 | 0.94 | 0.93 | 0.91 | 0.97 | 0.94 | 0.9 | 0.85 | 0.86 | 0.85 |
| 38 | 0.94 | 0.9 | 0.93 | 0.96 | 0.94 | 0.9 | 0.9 | 0.92 | 0.89 |
| 39 | 0.94 | 0.87 | 0.95 | 0.96 | 0.95 | 0.92 | 0.95 | 0.94 | 0.92 |
| 40 | 0.94 | 0.85 | 0.95 | 0.95 | 0.96 | 0.94 | 0.97 | 0.94 | 0.94 |
| 41 | 0.94 | 0.84 | 0.95 | 0.95 | 0.96 | 0.96 | 0.96 | 0.93 | 0.94 |
| 42 | 0.94 | 0.84 | 0.95 | 0.94 | 0.96 | 0.96 | 0.94 | 0.93 | 0.93 |
| 43 | 0.95 | 0.86 | 0.95 | 0.94 | 0.95 | 0.96 | 0.93 | 0.94 | 0.92 |
| 44 | 0.96 | 0.89 | 0.95 | 0.95 | 0.95 | 0.95 | 0.93 | 0.94 | 0.92 |
| 45 | 0.96 | 0.92 | 0.94 | 0.96 | 0.95 | 0.94 | 0.93 | 0.92 | 0.92 |
| 46 | 0.96 | 0.95 | 0.93 | 0.96 | 0.95 | 0.93 | 0.93 | 0.9 | 0.9 |
| 47 | 0.95 | 0.96 | 0.91 | 0.97 | 0.94 | 0.93 | 0.92 | 0.9 | 0.89 |
| 48 | 0.94 | 0.96 | 0.9 | 0.97 | 0.94 | 0.93 | 0.91 | 0.91 | 0.89 |
| 49 | 0.93 | 0.96 | 0.89 | 0.97 | 0.94 | 0.94 | 0.9 | 0.91 | 0.89 |
| 50 | 0.93 | 0.95 | 0.89 | 0.97 | 0.95 | 0.95 | 0.9 | 0.9 | 0.9 |
| 51 | 0.92 | 0.95 | 0.9 | 0.97 | 0.96 | 0.97 | 0.92 | 0.9 | 0.91 |
| 52 | 0.92 | 0.95 | 0.91 | 0.97 | 0.97 | 0.97 | 0.94 | 0.92 | 0.92 |
| 53 | 0.92 | 0.95 | 0.93 | 0.97 | 0.97 | 0.97 | 0.96 | 0.95 | 0.93 |
| 54 | 0.92 | 0.96 | 0.95 | 0.98 | 0.96 | 0.96 | 0.97 | 0.96 | 0.94 |
| 55 | 0.93 | 0.96 | 0.97 | 0.98 | 0.96 | 0.96 | 0.97 | 0.96 | 0.95 |
| 56 | 0.93 | 0.97 | 0.97 | 0.98 | 0.96 | 0.97 | 0.96 | 0.95 | 0.95 |
| 57 | 0.93 | 0.97 | 0.97 | 0.97 | 0.96 | 0.97 | 0.95 | 0.95 | 0.94 |
| 58 | 0.93 | 0.97 | 0.96 | 0.96 | 0.96 | 0.97 | 0.95 | 0.95 | 0.93 |
| 59 | 0.93 | 0.97 | 0.94 | 0.96 | 0.95 | 0.97 | 0.94 | 0.95 | 0.93 |
| 60 | 0.94 | 0.96 | 0.93 | 0.95 | 0.95 | 0.97 | 0.94 | 0.95 | 0.93 |
| 61 | 0.94 | 0.95 | 0.92 | 0.95 | 0.95 | 0.96 | 0.94 | 0.94 | 0.93 |
| 62 | 0.94 | 0.95 | 0.92 | 0.95 | 0.95 | 0.96 | 0.94 | 0.93 | 0.93 |
| 63 | 0.94 | 0.95 | 0.92 | 0.96 | 0.96 | 0.96 | 0.94 | 0.94 | 0.93 |
| 64 | 0.95 | 0.95 | 0.92 | 0.97 | 0.96 | 0.96 | 0.95 | 0.95 | 0.94 |
| 65 | 0.95 | 0.95 | 0.93 | 0.98 | 0.97 | 0.97 | 0.96 | 0.97 | 0.96 |
| 66 | 0.96 | 0.96 | 0.94 | 0.99 | 0.97 | 0.98 | 0.97 | 0.98 | 0.98 |
| 67 | 0.95 | 0.97 | 0.95 | 0.99 | 0.98 | 0.98 | 0.99 | 0.98 | 0.98 |
| 68 | 0.94 | 0.97 | 0.96 | 0.99 | 0.98 | 0.99 | 0.98 | 0.99 | 0.98 |
| 69 | 0.93 | 0.98 | 0.96 | 0.98 | 0.98 | 0.99 | 0.98 | 0.98 | 0.97 |
| 70 | 0.93 | 0.98 | 0.97 | 0.97 | 0.98 | 0.99 | 0.97 | 0.97 | 0.97 |
| 71 | 0.92 | 0.98 | 0.97 | 0.97 | 0.98 | 0.98 | 0.97 | 0.97 | 0.97 |
| 72 | 0.93 | 0.98 | 0.98 | 0.97 | 0.99 | 0.98 | 0.97 | 0.97 | 0.97 |
| 73 | 0.93 | 0.98 | 0.98 | 0.98 | 0.99 | 0.98 | 0.97 | 0.97 | 0.97 |
| 74 | 0.95 | 0.98 | 0.98 | 0.98 | 0.99 | 0.98 | 0.98 | 0.98 | 0.97 |
| 75 | 0.96 | 0.98 | 0.98 | 0.99 | 0.99 | 0.98 | 0.98 | 0.98 | 0.98 |
| 76 | 0.97 | 0.99 | 0.98 | 0.99 | 0.99 | 0.99 | 0.98 | 0.98 | 0.98 |
| 77 | 0.98 | 0.99 | 0.98 | 0.99 | 0.99 | 0.99 | 0.98 | 0.98 | 0.98 |
| 78 | 0.99 | 0.99 | 0.98 | 0.99 | 0.99 | 0.99 | 0.99 | 0.99 | 0.98 |
| 79 | 0.99 | 0.99 | 0.98 | 0.99 | 0.99 | 0.99 | 0.99 | 0.99 | 0.99 |
| 80 | 0.99 | 0.99 | 0.98 | 0.99 | 0.99 | 0.99 | 0.99 | 0.99 | 0.99 |
| 81 | 0.99 | 0.99 | 0.98 | 0.99 | 0.99 | 0.99 | 0.99 | 0.99 | 0.99 |
| 82 | 0.98 | 0.99 | 0.98 | 0.99 | 0.99 | 0.99 | 0.99 | 0.99 | 0.99 |
| 83 | 0.98 | 0.99 | 0.99 | 0.99 | 0.99 | 0.99 | 0.99 | 0.99 | 0.98 |
| 84 | 0.97 | 0.99 | 0.99 | 0.99 | 0.99 | 0.99 | 0.98 | 0.99 | 0.98 |
| 85 | 0.95 | 0.98 | 0.98 | 0.99 | 0.99 | 0.98 | 0.98 | 0.99 | 0.98 |
| 86 | 0.93 | 0.97 | 0.98 | 0.99 | 0.99 | 0.98 | 0.98 | 0.98 | 0.98 |
| 87 | 0.9 | 0.95 | 0.96 | 0.98 | 0.99 | 0.97 | 0.98 | 0.98 | 0.98 |
| 88 | 0.87 | 0.92 | 0.94 | 0.98 | 0.98 | 0.96 | 0.97 | 0.97 | 0.97 |
| 89 | 0.83 | 0.88 | 0.92 | 0.97 | 0.97 | 0.94 | 0.96 | 0.96 | 0.96 |

4. **Cell-specific Gene Expression Profiling**

Gene expression data were obtained from the Allen Human Brain Atlas (AHBA), based on postmortem human brains, providing comprehensive coverage of the adult brain (<http://www.brain-map.org>) (Hawrylycz et al. 2012). Isolated RNA was hybridized to custom 64K Agilent microarrays (58,692 probes) by the Allen Institute. Gene expression data for the left hemisphere were available for six donors. Gene-expression data from the AHBA were mapped to the Desikan-Killiany atlas (Desikan et al. 2006) in FreeSurfer space yielding up to 1269 labeled samples per brain inside or close to a FreeSurfer cortical region (French and Paus, 2015). The expression values of the mapped samples were mean averaged across microarray probes to provide a single expression value for each gene for a given sample. Median averages were used to summarize expression values within each cortical region and donor, which was followed by the median average across the six donors. This yielded a single value for each region representing the median profile for a given gene across the 34 cortical regions. We used previously developed scripts (Shin et al. 2018) to obtain gene expression profiles across the left hemisphere. Note that only genes (n=2511) with inter-regional profiles consistent across the six donors (AHBA) and across two datasets (AHBA and the BrainSpan Atlas) were considered in further analyses (Shin et al. 2018). Lists of genes expressed in specific cell types were obtained from Zeisel and colleagues (Zeisel et al. 2015) who obtained single-cell transcriptomes from 3005 cells from the somatosensory cortex (S1) and the CA1 hippocampus of mice. Gene expression was then clustered into nine classes each containing over 100 genes. Mouse genes were converted to human gene symbols with the “homologene” R package (Mancarci and French, 2019) The resulting classes of cell types and the number of marker genes in the consistent gene set were as follows: S1 pyramidal neurons (n = 73 human gene symbols), CA1 pyramidal neurons (n = 103), interneurons (n = 100), astrocytes (n = 54), microglia (n = 48), oligodendrocytes (n = 60), ependymal (n = 84), endothelial (n = 57), and mural (pericytes and vascular smooth muscle cells; n = 25).

**5.****Study Specific Gene Expression Database and Gene Co-expression Networks**

We generated a study-specific gene-expression database based on information from post-mortem cortical brain tissue from 572 unique donors, aged 0 and 102 years at the time of death, from five databases: the AHBA, the BrainCloud (Colantuoni et al. 2011; Jaffe et al. 2015), the Brain eQTL Almanac project (BrainEAC) (Trabzuni et al. 2011), the BrainSpan (http://brainspan.org) and the Genotype-Tissue Expression Project (GTEx) (GTEx Consortium, 2015). Gene expression was quantified using microarrays in AHBA, BrainCloud, and BrainEAC and RNA sequencing in BrainSpan and GTEx. Gene symbols in each database were updated, the gene expression values were log10 transformed, scaled within each cortical region. We then created a study specific database by pooling the values of the 15380 genes that were available in all five databases. This study-specific database included expression values of the majority (n=2321) of the consistent genes panel. For each database, gene symbols were updated, and their values were log10 transformed and scaled within each sampled region. Gene expression was then combined across all datasets. Only genes with expression values from all five databases were included in the final curated expression database (15380 genes). We tested the co-expression for all consistent genes (n = 2321; not all consistent genes were present in the 5 different datasets) using linear mixed models as implemented with *lme4* in R (Bates et al. 2015). The models were adjusted for age and sex with a subject identifier as a random intercept. Bound Optimization BY Quadratic Approximation (BOBYQA) (Powell, 2009) was set as optimizer parameter while the default options were used for the remaining parameters¨ For each consistent gene, the degree of co-expression was then determined by ranking the signed effect sizes of all linear models as implemented in the *t_to_eta2 function in effectsize* (Ben-Shachar et al. 2020)*.* This pipeline resulted in a 2321 × 15380 co-expression matrix.

**6. Permutation Analyses for the Correlation between Lifespan Thinning and Cell-Specific Gene Expression Profiles**

To assess the relationship between the cell-specific gene expression profiles and cortical thinning throughout the lifespan, we computed the Pearson’s correlation between each GAMM derivative estimated at 1-year shifts across the sample ((n = 87, one derivative for each chronological age from 4 to 89 years) and each consistent gene (n = 2511). Statistical significance established using resampling and permutation testing against a null distribution as per Vidal-Pineiro et al (2020). To achieve this, we generated pseudo-cell-specific panels, using a random selection of consistent genes of equal number to that of the each of the actual cell-specific gene panels. We calculated the Pearson’s correlation coefficient between each gene in these pseudo-panels and each cortical thinning profile and averaged the pseudo-coefficients. We then selected the mean pseudo-correlation with the maximum absolute value for each cortical thinning profile thus introducing Bonferroni-like correction for multiple comparisons at a within cell-type level while accounting for the non-independency of the thinning estimates. This process was repeated 1000 times to generate a null distribution. A two-sided p-value was computed as the proportion of peudo-correlations whose values exceeded the average for those in the actual cell-specific panel. We rejected the null hypothesis at P_FDR_ = 0.05 to control for comparisons for the nine cell-specific panels. Thus, statistical threshold was determined by combining Bonferroni-like correction for the number of cortical profiles and FDR correction the number of cell-specific panels.

**7. Code Availability**

The scripts for the analyses conducted will be available online at: https://github.com/AmirhosseinModabbernia/VirtualHistology/

B. **Supplementary Results**
1. **Cortical Thinning Estimates Across the Lifespan**

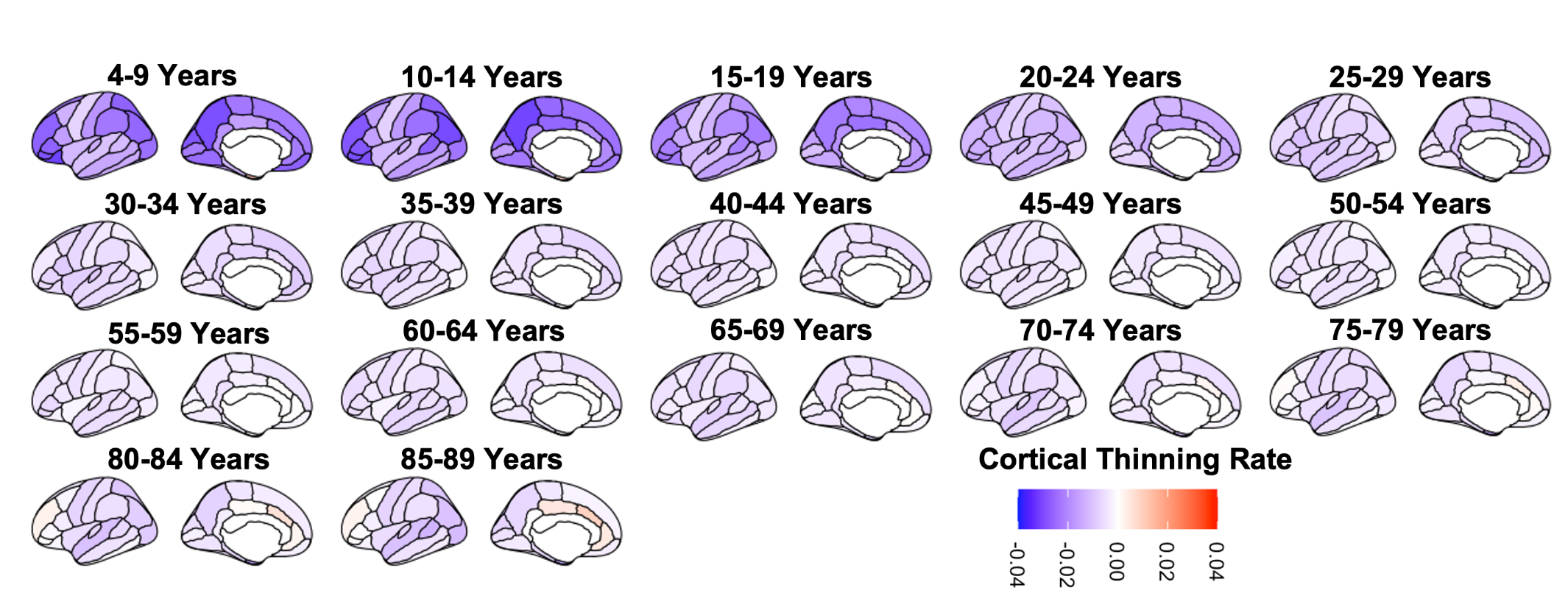
 **Supplementary Figure 2. Estimates of Cortical Thinning Rate Across the Lifespan**

2. **Sensitivity analyses of virtual histology to the Number of Knots in the General Additive Mixed Models for Cortical Thinning**

**Supplementary Figure 3. Effect of Varying Number of Knots in the Generalized Additive Mixed Models on Virtual Histology**

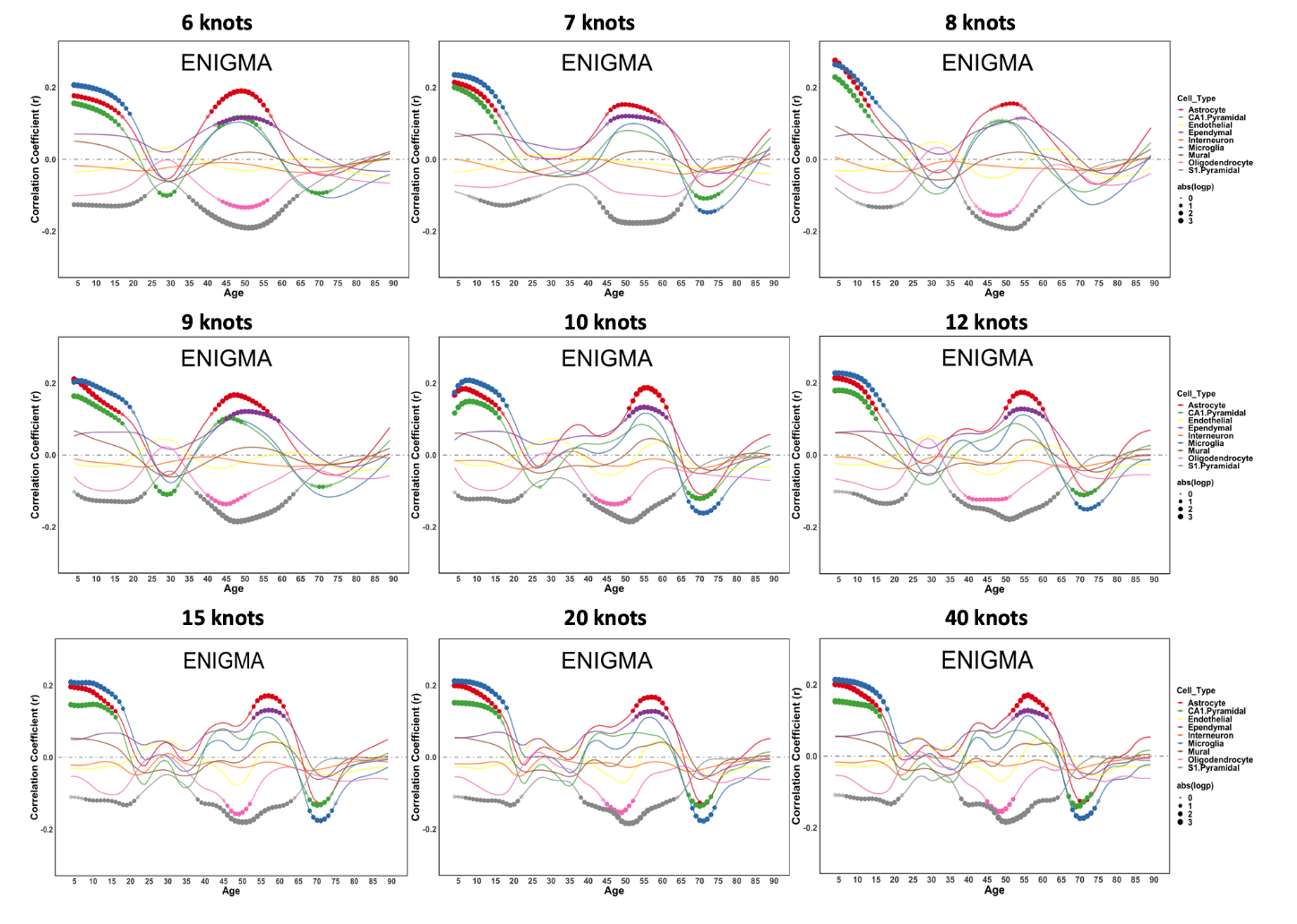

3. **Association of life-stage inter-cortical thinning profiles with gene co-expression and enrichment**

**Supplementary Figure 4. Cnetplot for the Linkages of Genes and Biological Processes in Early-Life**

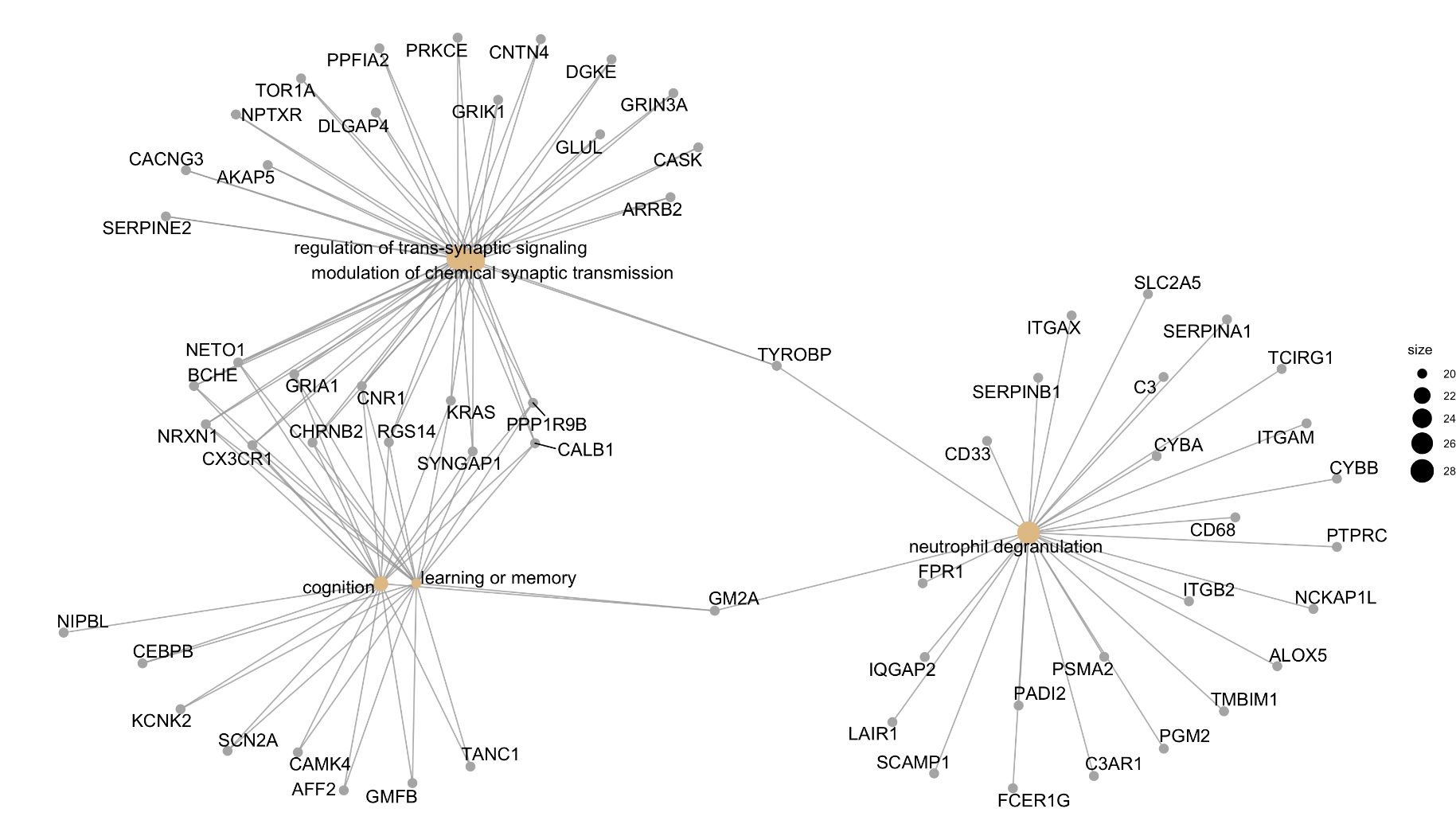

**Supplementary Figure 5. Cnetplot for the Linkages of Genes and Cellular Components in Early-Life**

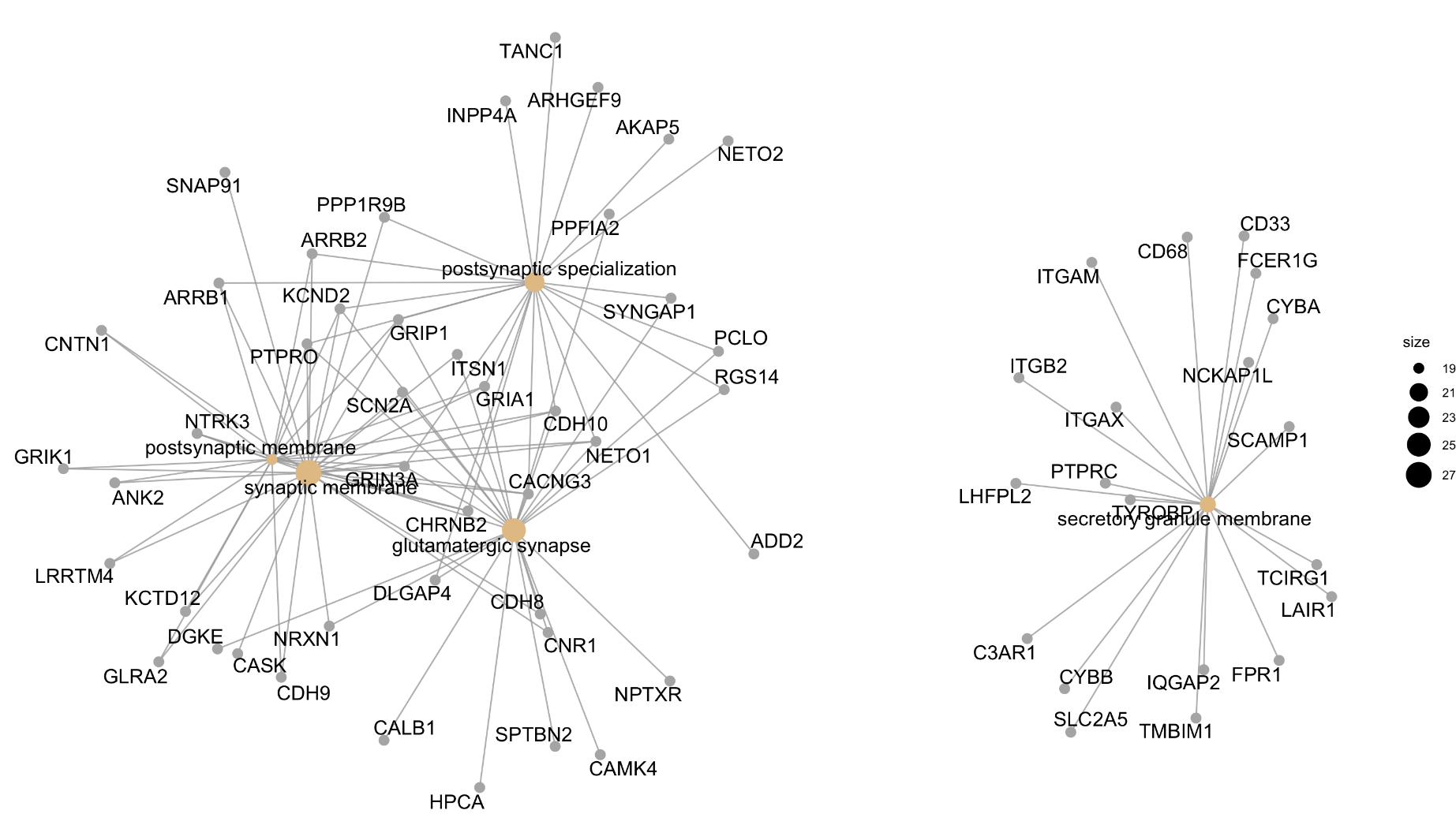

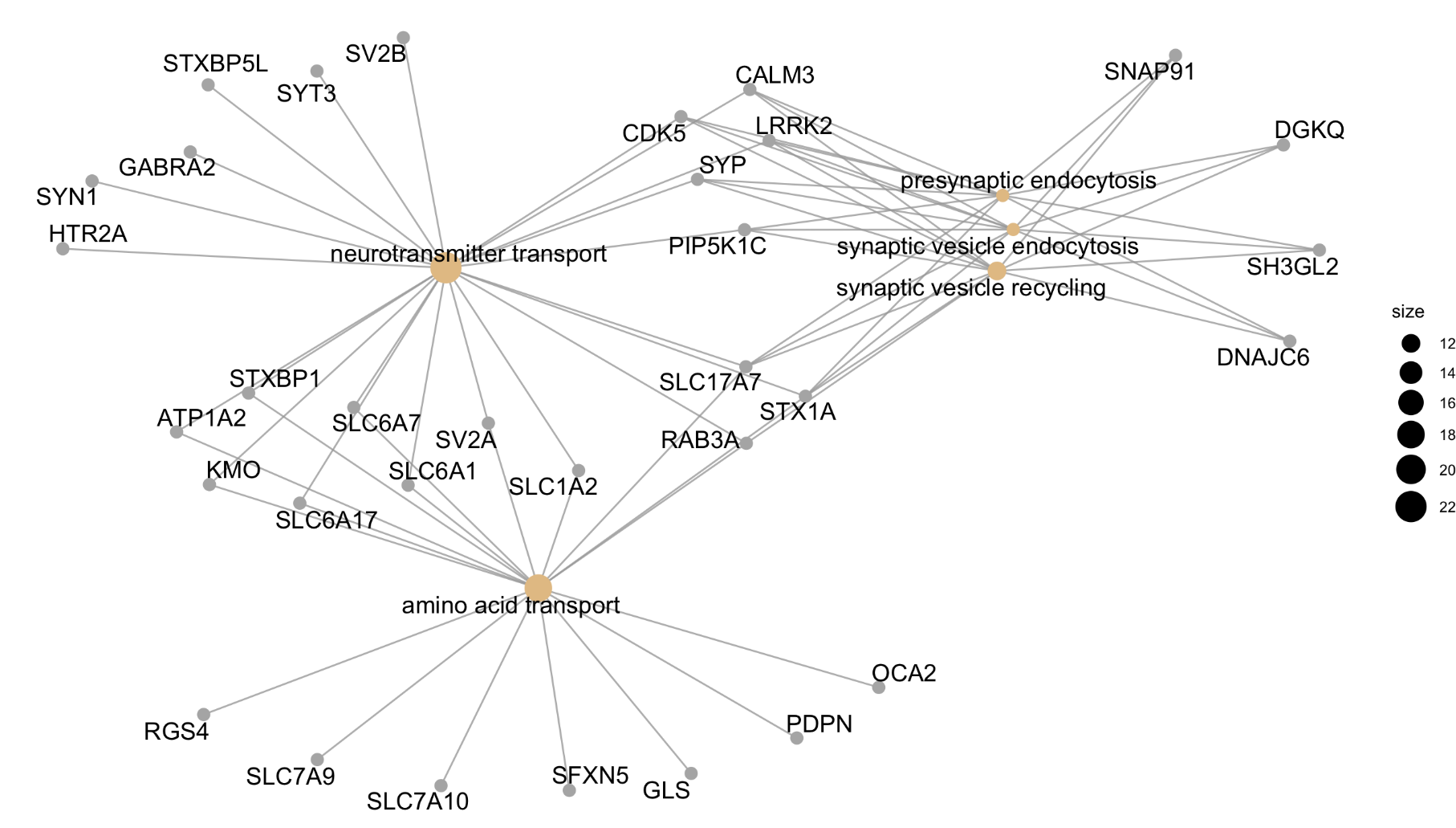
 **Supplementary Figure 6. Cnetplot for the Linkages of Genes and Biological Processes in Mid-Life**

**Supplementary Figure 7. Cnetplot for the Linkages of Genes and Cellular Processes in Mid-Life**

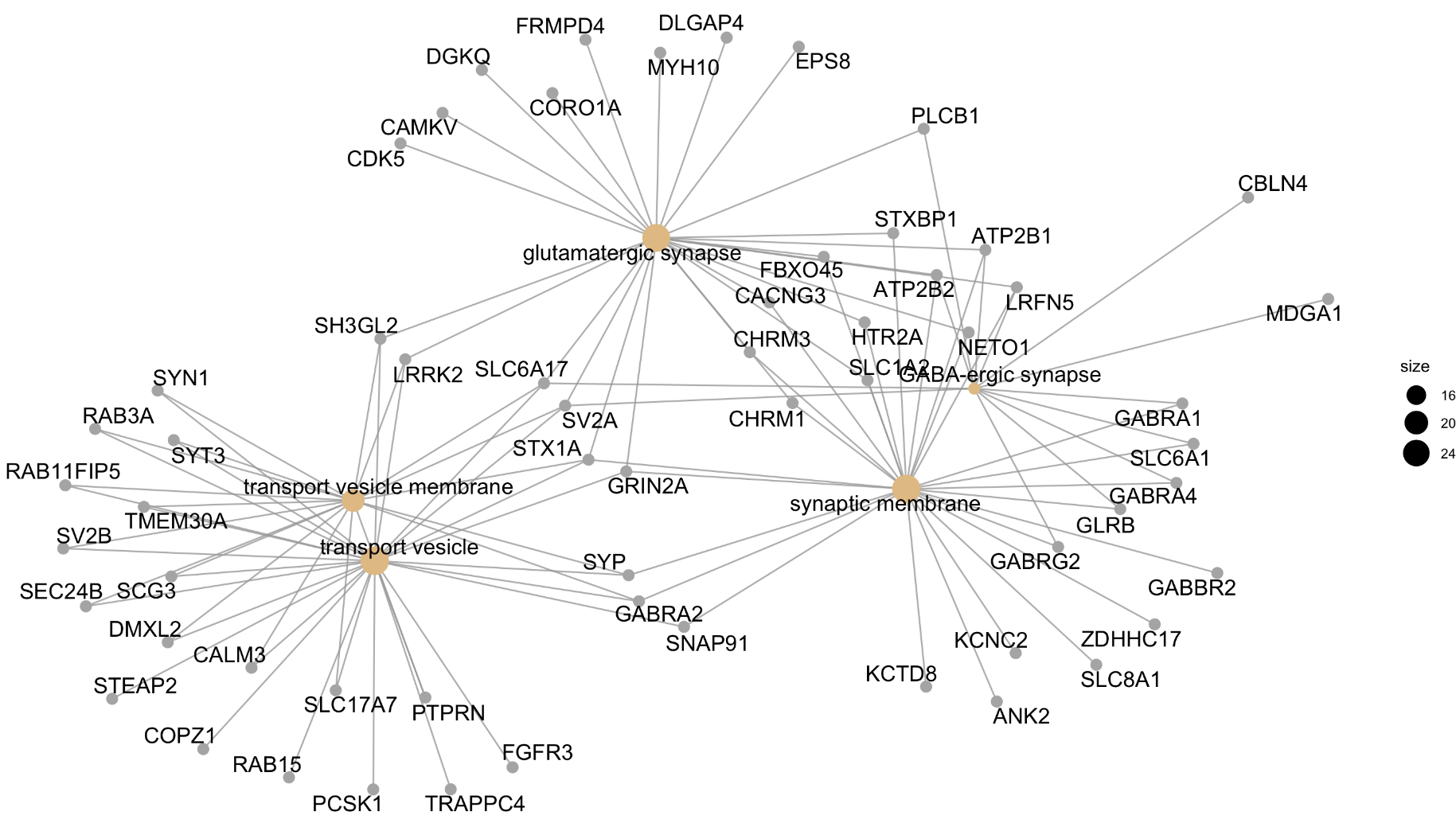

**Supplementary Figure 8. Cnetplot for the Linkages of Genes and Biological Processes in Late-Life**

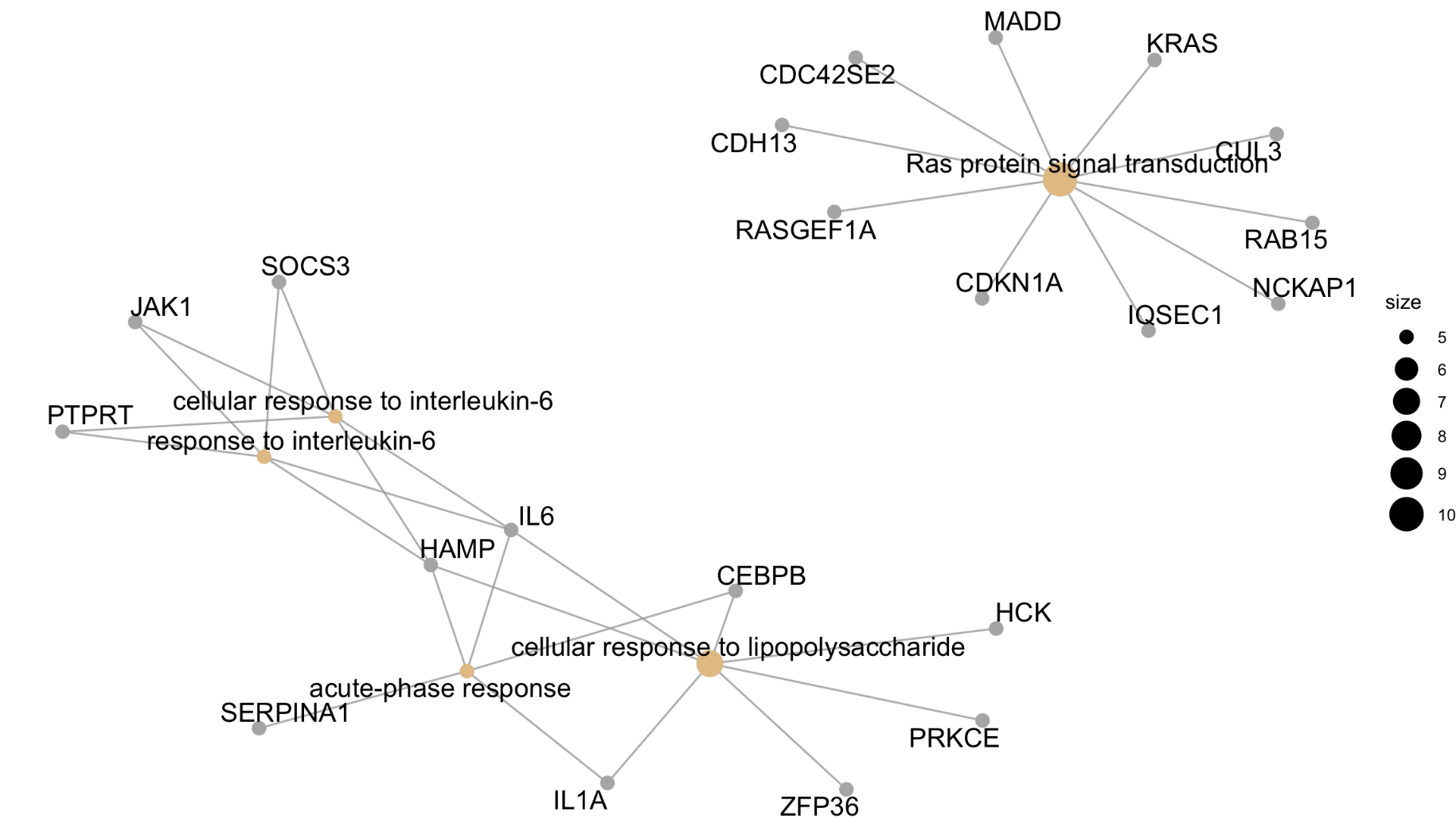

**Supplementary Figure 9. Cnetplot for the Linkages of Genes and Cellular Components in late-Life**

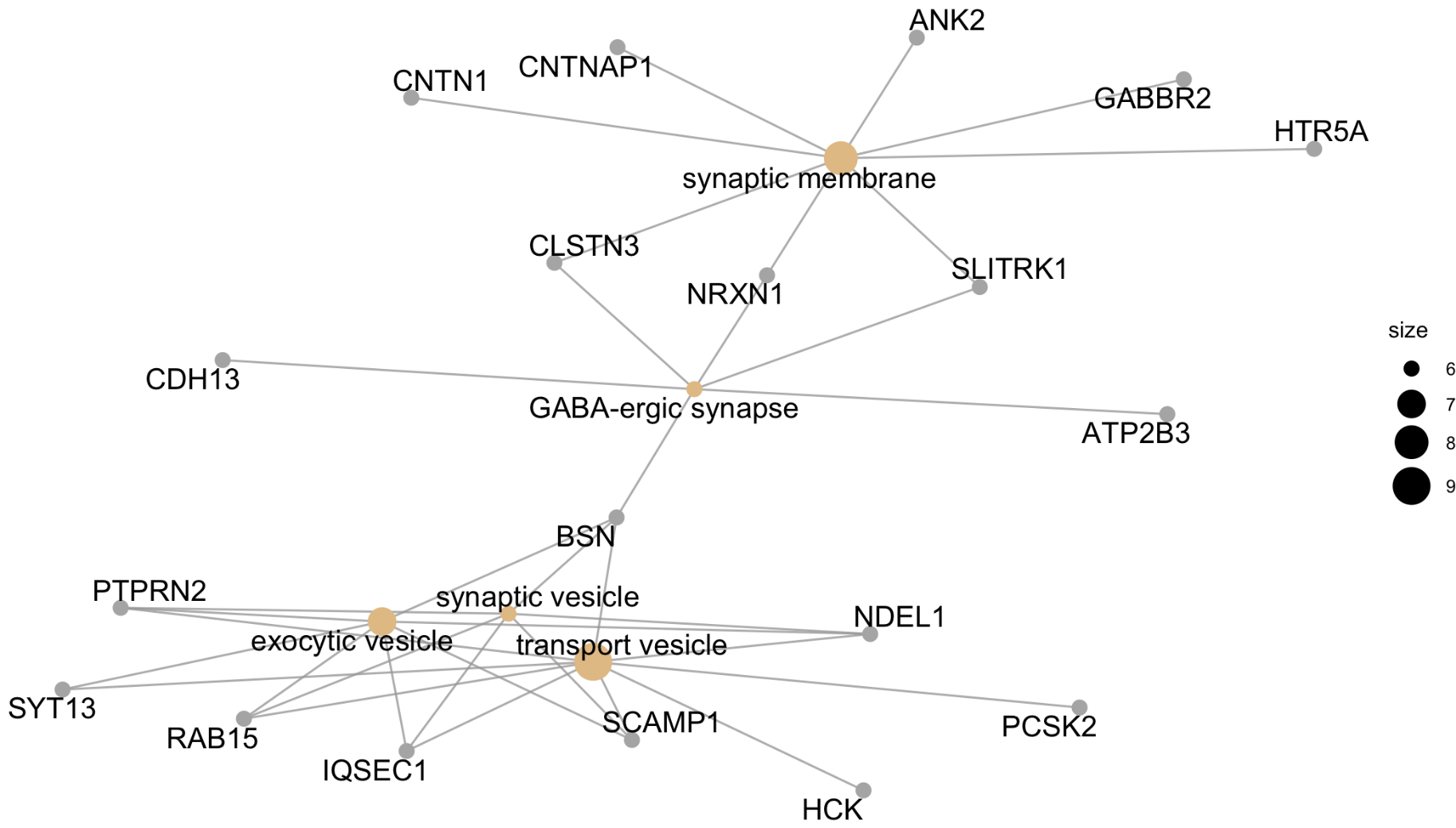

4. **Sensitivity Analyses for Gene Ontology**

Three sets of sensitivity analyses were performed to evaluate the robustness of gene ontology (GO) results. First, we repeated the GO analyses after systematically varying the percentage of the consistent genes from the top 1% to the top 10%, in 1% increments and percentage of co-expressed genes from 0.1% to 0.5% in 0.1% increments. Second, we repeated the GO analyses for each specific cell-type in each age group (i.e., early, mid-, late life). Third, we repeated the analyses for specific cell-types only in a period where their gene-expression profiles were significantly associated with cortical thinning.

Due to large number of the figures for the sensitivity analyses, these materials can be accessed online via

<https://doi.org/10.6084/m9.figshare.17099789>

<https://doi.org/10.6084/m9.figshare.17100035>

<https://doi.org/10.6084/m9.figshare.17099786>

**C.Supplementary References**

Bates D, Mächler M, Bolker B, Walker S. 2015. Fitting linear mixed-effects models using lme4. Journal of Statistical Software, 67: 1-48

Ben-Shachar MS, Lüdecke D, Makowski D. 2020. effectsize: Estimation of effect size indices and standardized parameters. *Open Source Softw*. 5:2815.

BrainSpan Atlas of the Developing Human Brain. <http://brainspan.org>

Colantuoni C, Lipska BK, Ye T, Hyde TM, Tao R, Leek JT, Colantuoni EA, Elkahloun AG, Herman MM, Weinberger DR, Kleinman JE. 2011. Temporal dynamics and genetic control of transcription in the human prefrontal cortex. *Nature*. 478:519-523

Desikan RS, Ségonne F, Fischl B, Quinn BT, Dickerson BC, Blacker D, Buckner RL, Dale AM, Maguire RP, Hyman BT et al. 2006. *Neuroimage*. 31:968-980.

French L, Paus T. 2015. A FreeSurfer view of the cortical transcriptome generated from the Allen Human Brain Atlas. *Front Neurosci*. 9:323.

GTEx Consortium. 2015. Human genomics. The Genotype-Tissue Expression (GTEx) pilot analysis: multitissue gene regulation in humans. *Science*. 348:648-660.

Hawrylycz MJ, Lein ES, Guillozet-Bongaarts AL, Shen EH, Ng L, Miller JA, van de Lagemaat LN, Smith KA, Ebbert A, Riley ZL et al. 2012. An anatomically comprehensive atlas of the adult human brain transcriptome. *Nature*. 489:391-399.

Jaffe AE, Hyde T, Kleinman J, Weinbergern DR, Chenoweth JG, McKay RD, Leek JT, Colantuoni C. 2015. Practical impacts of genomic data “cleaning” on biological discovery using surrogate variable analysis. *BMC Bioinformatics*. 16:372.

Mancarci O, French L. 2019. Homologene: quick access to homologene and gene annotation updates.

<https://cran.r-project.org/web/packages/homologene/homologene.pdf>

Powell MJD. 2009, "The BOBYQA algorithm for bound constrained optimization without derivatives", Report DAMTP 2009/NA06, University of Cambridge.

Shin J, French L, Xu T, Leonard G, Perron M, Pike GB, Richer L, Veillette S, Pausova Z, Paus T. 2018. Cell-Specific Gene-Expression Profiles and Cortical Thickness in the Human Brain. *Cereb Cortex*. 28:3267-3277.

Trabzuni D, Ryten M, Walker R, Smith C, Imran S, Ramasamy A, Weale ME, Hardy J. 2011. Quality control parameters on a large dataset of regionally dissected human control brains for whole genome expression studies. *J Neurochem*. 119:275-282.

Vidal-Pineiro D, Parker N, Shin J, French L, Grydeland H, Jackowski AP, Mowinckel AM, Patel Y, Pausova Z, Salum G et al. 2020. Cellular correlates of cortical thinning throughout the lifespan. Sci Rep. 10:21803.

Zeisel A, Muñoz-Manchado AB, Codeluppi S, Lönnerberg P, La Manno G, Juréus A, Marques S, Munguba H, He L, Betsholtz C et al. 2015. Brain structure. Cell types in the mouse cortex and hippocampus revealed by single-cell RNA-seq. *Science*. 347:1138-1142.
